# Stereoselectivity and functional plasticity of a common ligand-binding pocket in TRPM3

**DOI:** 10.1101/2025.08.27.671268

**Authors:** Bahar Bazeli, Alexander V Shkumatov, Stephan Schenck, Jean-Christophe Vanherck, Annelies Janssens, Robbe Roelens, Joris Vriens, Thomas Voets, Janine D Brunner

## Abstract

The transient receptor potential melastatin 3 (TRPM3) channel is a key mediator of peripheral pain signaling, and pathogenic mutations in TRPM3 are linked to neurodevelopmental delay and epilepsy. Despite the therapeutic promise of TRPM3 modulators, the molecular mechanisms by which ligands modulate channel gating remain poorly understood. Here, we combine cryo-electron microscopy (cryo-EM) with functional analyses to characterize a promiscuous ligand-binding pocket formed by transmembrane helices S1–S4. This pocket accommodates several chemically diverse plant-derived and synthetic agonists and antagonists. We uncover unanticipated stereoselectivity of TRPM3 for the non-natural enantiomer of the flavonoid antagonist isosakuranetin and the *(R)*-enantiomer of the synthetic agonist CIM0216. Mutations within this pocket—including newly identified patient variants—alter ligand affinity and, in some cases, invert the functional outcome of ligand binding. These findings reveal the stereoselectivity and functional plasticity of the TRPM3 ligand-binding pocket, highlighting how subtle changes in the molecular interactions can produce divergent effects on channel gating, with important ramifications for TRPM3-targeted drug development and therapy.

---

Transient Receptor Potential melastatin 3 (TRPM3) is a temperature-sensitive ionotropic neurosteroid receptor within the TRP superfamily of cation channels ^1,2^. In the peripheral nervous system, TRPM3 is expressed in a subset of somatosensory neurons from the dorsal root and trigeminal ganglia, where the channel contributes to the detection of noxious heat ^2–5^. Under pathological conditions such as inflammation or neuropathy, TRPM3 channel activity is increased, contributing to sensory hypersensitivity and ongoing pain ^3,4,6–10^. In addition, TRPM3 is expressed in various neuronal and non-neuronal cells in the developing and adult brain, but its role in brain function is not yet well established ^11,12^. Notably, gain-of-function mutations in TRPM3 cause a spectrum of neurodevelopmental disorders, characterized by a variety of neurological symptoms, including developmental delay, hypotonia, seizures and often alterations in heat- or pain sensitivity ^11,13–19^. Therefore, strategies to modify TRPM3 activity represent important new therapeutic avenues to treat pathological pain and brain disorders.

Currently, selective TRPM3 modulators approved for use in humans are not yet available. Nevertheless, earlier research has demonstrated that TRPM3 is inhibited by naturally occurring polyphenoles such as the flavonoid isosakuranetin and the deoxybenzoin compound ononetin, and by the antiseizure drug barbiturate congener primidone ^20–22^. In preclinical studies, these compounds have shown efficacy in reducing nociceptive behavior in rodent models of inflammatory and neuropathic pain, at least partly through inhibition of TRPM3 activity ^3,4,6,9,10,20,22^. Moreover, primidone suppressed electroencephalographic abnormalities and resulted in significant developmental improvement in a small-scale clinical study with patients suffering from neurodevelopmental disease due to TRPM3 gain-of-function mutations ^19,23^. However, primidone and its metabolite phenobarbital also have sedative effects, thereby limiting its use in patients ^19^. A better understanding of the molecular mechanisms of TRPM3 channel modulation by small molecules may form the basis of the rational development of more selective TRPM3-targeting drugs. In this regard, recent studies have already provided important insights into the overall structure of TRPM3, and highlighted binding sites of the small-molecule inhibitor primidone, and the agonists pregnenolone sulphate (PS) and CIM0216 ^24,25^. Yet, many important questions remain unanswered regarding the exact interaction mode of these and other small molecules with TRPM3, and how this affects channel gating.

In this study, we present cryo-EM structures of TRPM3 in the apo-state, and in complex with the antagonists isosakuranetin, ononetin, primidone, and with the agonist CIM0216. Supported by functional analyses of structure-guided mutants, we establish that these ligands have overlapping interaction poses in a cavity formed by S1-S4. Notably, we found that the binding pocket displays a strong stereoselectivity for isosakuranetin and CIM0216. We chirally resolved the commonly used racemic mixtures of these compounds, and demonstrate strongly diverging and even opposing effects for the respective *(R)*- and *(S)*-enantiomers. Moreover, we identified residues in the ligand binding pocket which, upon mutagenesis, affect the functional outcome of ligand binding. Further, our primidone- and CIM0216-bound structures reveal relevant differences for the interaction of the ligands with the pocket-lining residues in comparison to a previously published study. Finally, we report two patients with neurodevelopmental disorder carrying two distinct mutations at residues lining the binding pocket, which lead to gain of channel function combined with severely altered responses to primidone and other ligands. Our findings indicate that the S1-S4 cavity in TRPM3 represents a functionally plastic ligand binding site, where subtle changes in ligand-channel interaction can lead to opposing effects on channel gating. These results have important ramifications for the development of TRPM3-targeting small-molecules for therapeutic applications, and in particular for the treatment of patients with disease-causing variants in this region.

## Results

### Structures of TRPM3 in the apo state and in complex with antagonists

To obtain insights into how TRPM3 antagonists interact with the channel, we performed cryo-EM single particle analysis of mouse TRPM3 in the absence of ligands and in the presence of isosakuranetin, ononetin or primidone. To this end, we purified a truncated form of mouse TrpM3 isoform ±2 (amino acids 1-1344) from a stable HEK cell line and incubated the protein with the respective antagonists for 15-30 min prior to plunge-freezing. The final 3D reconstructions were resolved to 3.20 Å in the case of TrpM3 in the APO-state (TRPM3_APO_), and 2.30, 2.61 and 3.28 Å for TrpM3 bound to isosakuranetin (TRPM_ISO_), ononetin (TRPM3_ONO_) and primidone (TRPM3_PRM_), respectively (Figure 1A-C, Supplementary Figure 1-4, Supplementary Table 1). Generally, compared to the cytoplasmic N- and C-terminal densities the transmembrane (TM) part which is the focus of this study is resolved to a higher resolution, with local resolutions of up to 2Å (Supplementary Figure 3). The overall architecture adopts a fold that is typical for the TRPM family: the cytosolic portion is built of a substantial N-terminal melastatin homology region (MHR1-4) and a C-terminal coiled-coil domain while the transmembrane domain (TM) is formed through 6 helices of which S1-S4 bundle as voltage-sensor-like-domain (VSLD) and the domain-swapped S5 and S6 helices establish the pore (Supplementary Figure 5A,B). The atomic models adopt overall a very similar tetrameric assembly and align well with each other as well as with a previously published structure (8ED7) ^24^ with r.m.s.d. values of less than 1 Å (1.4 Å for 9B29 ^25^). According to a HOLE analysis the conformations in our models represent non-conductive states showing close apposition for G1066 in the selectivity filter (SF) and I1121 and N1125 at the lower gate, respectively (note that for referring to specific amino acids in this study we used the human consensus numbering^11^; Supplementary Figure 5A-D, Supplementary Table 2). Since we did not add any reducing agents to our preparations, we observe a nearly complete cap domain including the formation of a disulfide bridge between residues C1077 and C1094 (Supplementary Figure 5E).

**Figure 1:**
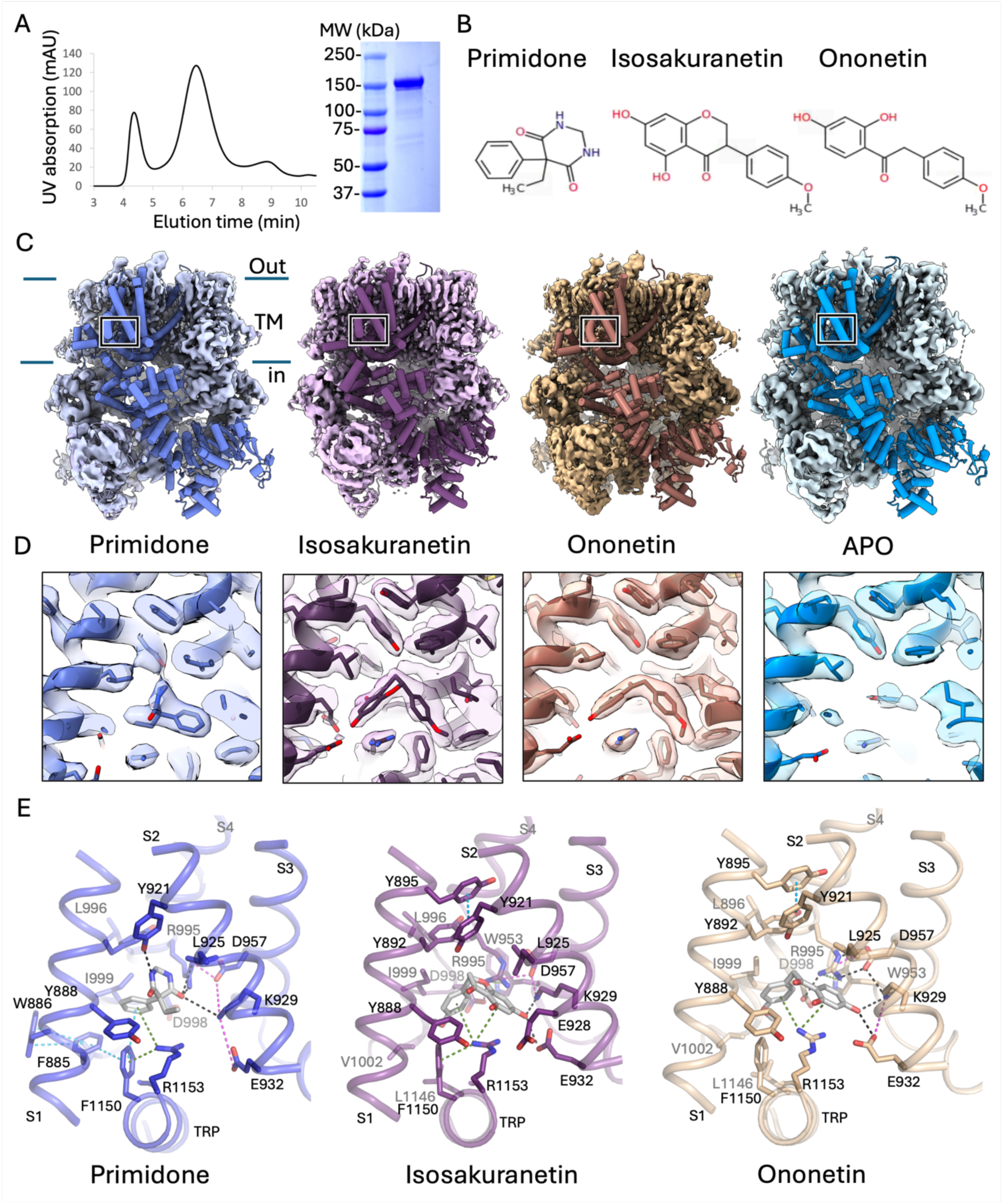
3D reconstructions and atomic models of TRPM3APO and of TRPM3 in the presence of antagonists. a) Size exclusion chromatography of TRPM3 (left) and SDS-PAGE analysis of the peak fraction. b) Chemical drawings of primidone, isosakuranetin and ononetin. c) Overall structures of TRPM3 bound to primidone (violet blue), isosakuranetin (violet), ononetin (wheat) and in the apo-state (marine) displayed as surface representation with one subunit shown as cylinders. Contouration in ChimeraX is at 0.039, 0.062, 0.061 and 0.113 σ for primidone, isosakuranetin, ononetin and the apo-state, respectively. d) Close-up view from the side on the VSLD ligand binding pocket with ligands and interacting residues lining S1-S4 shown as sticks. e) Close-up views on the ligand binding sites are shown with important interactions highlighted as dashed lines; hydrogen bonds in black; salt bridges in pink; cation-π stacks in green and π-π stacks in cyan. Hydrophobic contacts are not specifically marked. In (D) and (E) colours are as in (C).

For the three structures obtained in the presence of the different antagonists, we observed clear densities in a cavity in the VSLD surrounded by amino acid residues from S1-S4 and the C-terminal TRP domain, which were absent in TRPM3_APO_ (Figure 1D). The quality of the cryo-EM densities allowed unambiguous model building of the interactions between isosakuranetin, ononetin and primidone with the channel. In the case of primidone, the obtained binding mode of the ligand differs substantially from a recently published structure (PDB 9B28) ^25^. Primidone is a pyrimidone consisting of a dihydropyrimidine-4,6(1H,5H)-dione (heterocycle) substituted by an ethyl and a phenyl group at position 5. Compared to 9B28, primidone is tilted by ∼45° along an axis parallel to the phenyl and ethyl groups and moved about 1Å towards the extracellular side. This tilting and displacement of primidone results in a different set of TRPM3 residues interacting with primidone and diverging types of interactions (Supplementary Figure 6A-C). The side chains of Y921 (S2), K929 (S2) and R995 (S4) form hydrogen bonds with an amine and a carbonyl oxygen of the primidone heterocycle ring. R995 and K929 are further positioned through two salt bridge interactions with D957 and E932 on S2 and a hydrogen bond of R995 with D998 on S4, respectively (Figure 1E, left). The guanidinium group of R1153 of the TRP domain engages in a hydrophobic interaction with the ethyl group of the primidone heterocycle, in an interaction geometry that is reminiscent of arginine interacting with hydrophobic amino acid side chains ^26^. Arg1153 is additionally involved in cation-π interactions with the phenyl moiety of primidone as well as with F1150 of the TRP domain making this residue central for the coordination of the primidone ligand (Figure 1E, Supplementary Figure 6A). Phe1150 takes part in a T-shaped π stacking with the phenyl group of primidone and additionally with F885 of S1 which itself provides a hydrophobic contact with primidone and is stabilized through π-stacking with W886. Further hydrophobic interactions with primidone are mediated through the closely positioned Y888, L925 and I999 on S1, S2 and S4, respectively.

Different from the published structure of 9B28 we note additional (previously not reported) interactions with primidone, including hydrogen bonds between with K929 (S2) and R995 (S4) as well as a cation-π stack with R1153 (Figure 1E, Supplementary Figure 6A). The reported (pseudo-) symmetrical coordination (in particular through the arginines 995 and 1153 with the carbonyl-oxygens in primidone) presented in a recent study cannot be reconciled in our structure. Further, upon closer inspection during a reanalysis (using pymol, the Protein Ligand Interaction Profiler (PLIP) and LigPlot+) of the corresponding PDB entry 9B28 we cannot substantiate the previously presented network of interactions of TRPM3 with primidone (Supplementary Figure 6A,D, 7A). In 9B28 only two direct interactions occur according to our reanalysis; the π-π stack of F1150 with the phenyl ring of primidone like in the structure described herein, and a hydrogen bond between Y921 and the pyrimidinedione, that is in this case mediated through the other heterocycle amine group, due to the different positioning of primidone inside the VSLD binding pocket in the two structures. Supplementary Figure 6A presents direct interactions with primidone identified in 9B28 during our reanalysis. The positioning of the primidone molecule and the obtained network of interactions in the ligand binding pocket is thus significantly different between previously published work and our primidone-bound structure.

The binding sites of isosakuranetin and ononetin, herein described for the first time, overlap substantially with that of primidone (Figure 1C-E, Supplementary Figure 8A-C). Ononetin represents a natural product found in the spiny restharrow *Ononis spinosa* (Fabaceae), a plant that has been widely used for medicinal purposes, e.g. also as an anti-inflammatory analgesic in Russian traditional medicine ^27^. Within the deoxybenzoin chemical group, ononetin is an aryl ketone, with a 1-(2,4-dihydroxyphenyl)-2-(4-methoxyphenyl)ethanone structure. It is also structurally related to flavanones, a subgroup of flavonoids, where Isosakuranetin is a members of. Isosakuranetin is a naturally occurring O-methylated dihydroxy flavonoid, 5,7-dihydroxy-4’-methoxyflavanone, that is found in various plant sources, for example citrus fruits. Isosakuranetin and Ononetin interact in very similar manner with TRPM3, in accordance with their structural relationship (Figure 1B-D). Both antagonists, through their dihydroxybenzene moiety, make hydrogen bonds with the side chains of K929 and E932 on S2 which is part of a highly stabilized network of salt bridges, hydrogen bonds and cation-π stacks involving the side chains of E932 (S2), D957 and W953 (S3), R995 and D998 (S4). R1153 of the TRP domain is involved in cation-π stacks with the dihydroxybenzene and methoxybenzene groups and with F1150, strengthening the interaction network (Figure 1E, Supplementary Figure 7B,C). Y888 and Y892 (S1), Y921, and L925 (S2), I999 and V1002 (S4) as well as L1146 and F1150 (TRP) are in close proximity and mediate hydrophobic interactions with both antagonists.

### Stereoselectivity of isosakuranetin antagonism

Isosakuranetin is a natural compound produced in the flavonoid biosynthetic pathway in plants, containing a single chiral center at the C-2 carbon of the flavanone backbone ^28^. In the biosynthetic pathway, the enzyme chalcone isomerase catalyzes the intramolecular cyclization of chalcones to flavanones, which defines the stereochemistry of the C-2 chiral center, yielding a 100,000:1 preference for the synthesis of *(2S)*-flavanones over *(2R)*-flavanones ^29,30^. Accordingly, in publications describing its effects on TRPM3 as well as in curated databases such as the IUPHAR/BPS guide to pharmacology, isosakuranetin is consistently depicted as the *(S)-*enantiomer (IUPAC name: *(2S)*-5,7-dihydroxy-2-(4-methoxyphenyl)-2,3-dihydrochromen-4-one). When scrutinizing the cryo-EM-data obtained in the presence of commercially available, plant-derived isosakuranetin, we were surprised to see that the high-resolution EM density in the binding pocket corresponded to the non-natural *(R)-*enantiomer as judged from the quality of the density and from the fitting statistics (Figure 2A). We therefore examined the chiral purity of several batches of commercial isosakuranetin using chiral HPLC, and consistently observed two peaks corresponding to the *(R)-* and (S)-enantiomer, respectively, indicating that the product represents a racemic mixture (Figure 2B). Following purification, we compared the antagonist efficacy of the original, racemic isosakuranetin with that of the two separated enantiomers using a jRCAMP1b-based calcium imaging assay (Figure 2C). In this assay, cells were pre-incubated with different concentrations of racemic isosakuranetin or its enantiomers, followed by a test stimulus using PS (50 μM). Finally, cells were exposed to the ionophore ionomycin in the presence of excess extracellular Ca^2+^ (20 mM), to obtain saturation of the jRCAMP1b signal for normalisation (Figure 2C). Note that non-transfected HEK293 cells did not show any detectable changes in jRCAMP1b fluorescence in the presence of any of the TRPM3 ligands tested in this study (Supplementary Figure 9). For racemic isosakuranetin we found it to inhibit TRPM3 with an IC_50_ value of ∼1μM. Surprisingly, the naturally produced *(S)*-enantiomer, generally considered to be a potent TRPM3 antagonist, was largely ineffective, with less than 20% inhibition at the highest tested concentration (100 μM). In contrast, potent inhibition was obtained with the non-natural *(R)-*enantiomer, with an IC_50_ of ∼0.5μM. Taken together, these results indicate that commercially available, plant-derived isosakuranetin represents a racemic mixture, and that, contrary to what was generally assumed, the *(R)-* but not the (S)-enantiomer is responsible for the antagonistic effect at TRPM3. These measurements, together with the clearly resolvable *(R)-*isosakuranetin in our cryo-EM structure (Figure 2), provide compelling evidence for the specific interaction of this non-natural enantiomer with the channel.

**Figure 2:**
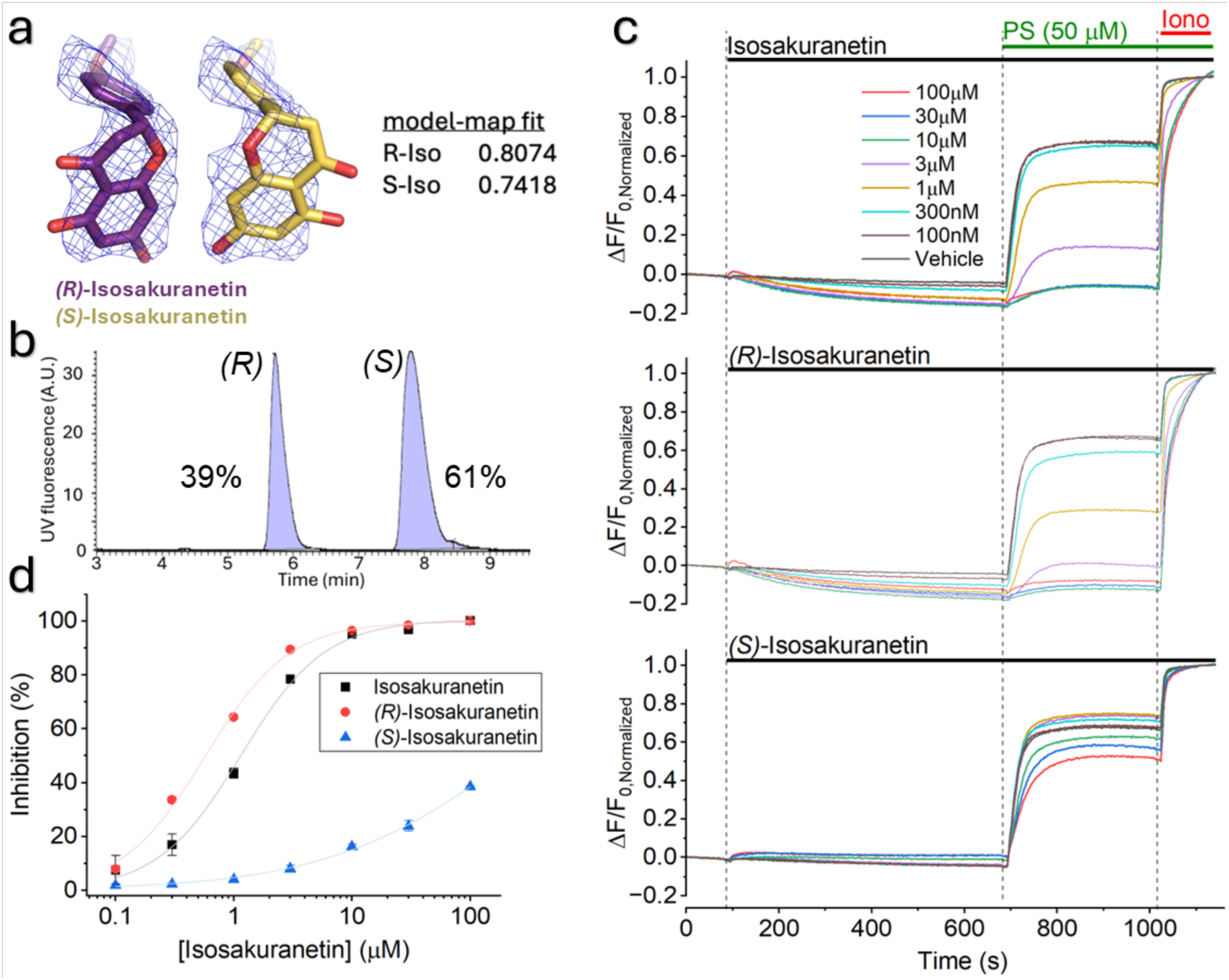
The non-natural *(R)*-isosakuranetin – not the natural *(S)*-isosakuranetin - is a potent TRPM3 antagonist. a) Cryo-EM density at the ligand binding site for isosakuranetin. R-isosakuranetin is shown in violet-purple and S-isosakuranetin in yellow. The density is contoured at a level of 8σ in pymol and the model-map fit values as calculated in ChimeraX is provided at the bottom. b) Chiral HPLC showing two peaks in commercial isosakuranetin. c) Normalized fluorescence traces showing changes in jRCAMP1b fluorescence in cells expressing WT human TRPM3, showing the effect of application of various concentrations of racemic isosakuranetin and its two enantiomers on responses evoked by PS. At the end of the experiment, a saturating ionomycin response was evoked for normalisation. (C) Concentration-response curves showing the inhibition of the PS-evoked response by racemic isosakuranetin and its two enantiomers. d) Concentration-inhibition curves for racemic isosakuranetin and the two enantiomers.

### Stereoselectivity of CIM0216 agonism and potentiation

CIM0216 (2-(3,4-Dihydroquinolin-1(2*H*)-yl)-*N*-(5-methyl-1,2-oxazol-3-yl)-2-phenylacetamide) potently activates TRPM3 and also acts as a potentiator of PS-induced channel activation. In a recent study, based on an incompletely resolved EM density and molecular dynamics simulations, a binding pose for CIM0216 was proposed in the S1-S4 pocket, partly overlapping with the binding site for primidone ^25^. It is important to note that CIM0216 used in the literature, including the study by Yin et al. ^25^, is a racemic mixture of *(R)-*CIM0216 and *(S)*-CIM0216. The (relative) potency of the two enantiomers has not been reported, nor is it known whether only one or both enantiomers contributed to the EM density detected in this previous work ^25^. To address this, we prepared pure *(R)-*CIM0216 and *(S)*-CIM0216, and tested the two enantiomers along with the racemic CIM0216 for their agonistic activity on TRPM3 using the jRCAMP1b-based calcium imaging assay. These experiments revealed that *(R)-* CIM0216 is a potent and full agonist of TRPM3, with an EC_50_ of ∼530 ± 14 nM, whereas (S)-CIM0216 is much less potent, showing detectable responses only at concentrations ≥3 μM (Figure 3A,B; Supplementary Figure 9). Racemic CIM0216 also evoked robust responses starting already at sub-micromolar concentrations, and with an EC_50_ value about 2-fold higher than for *(R)-*CIM0216 (1.33 ± 0.13 μM). To evaluate the potentiating effect of the different CIM0216 enantiomers, application of the compounds at different concentrations in the jRCAMP1b-based calcium imaging assay was followed by an invariant stimulation with PS (50 μM), and the enhancement of the amplitude of the final increase in fluorescence compared to vehicle was calculated. At concentrations that did not evoke a detectable response, racemic CIM0216 as well as the two enantiomers potentiated the response to PS, but with different potency. *(R)-*CIM0216 enhanced PS responses with an EC_50_ of 46 ± 5 nM, whereas *(S)*-CIM0216 was much less potent, with an EC_50_ of 710 ± 120 nM (Figure 3A,C). For racemic CIM0216 we obtained an intermediate EC_50_ of 75 ± 10 nM. Overall, these findings indicate that both the agonist and the potentiator activity of CIM0216 are primarily mediated by *(R)-*CIM0216, suggesting that this enantiomer may have a higher affinity for the binding pocket. The binding pose proposed in recent structural work ^25^, which for undisclosed reasons was modelled using the *(S)*-CIM0216 enantiomer, therefore needed to be revisited.

**Figure 3.**
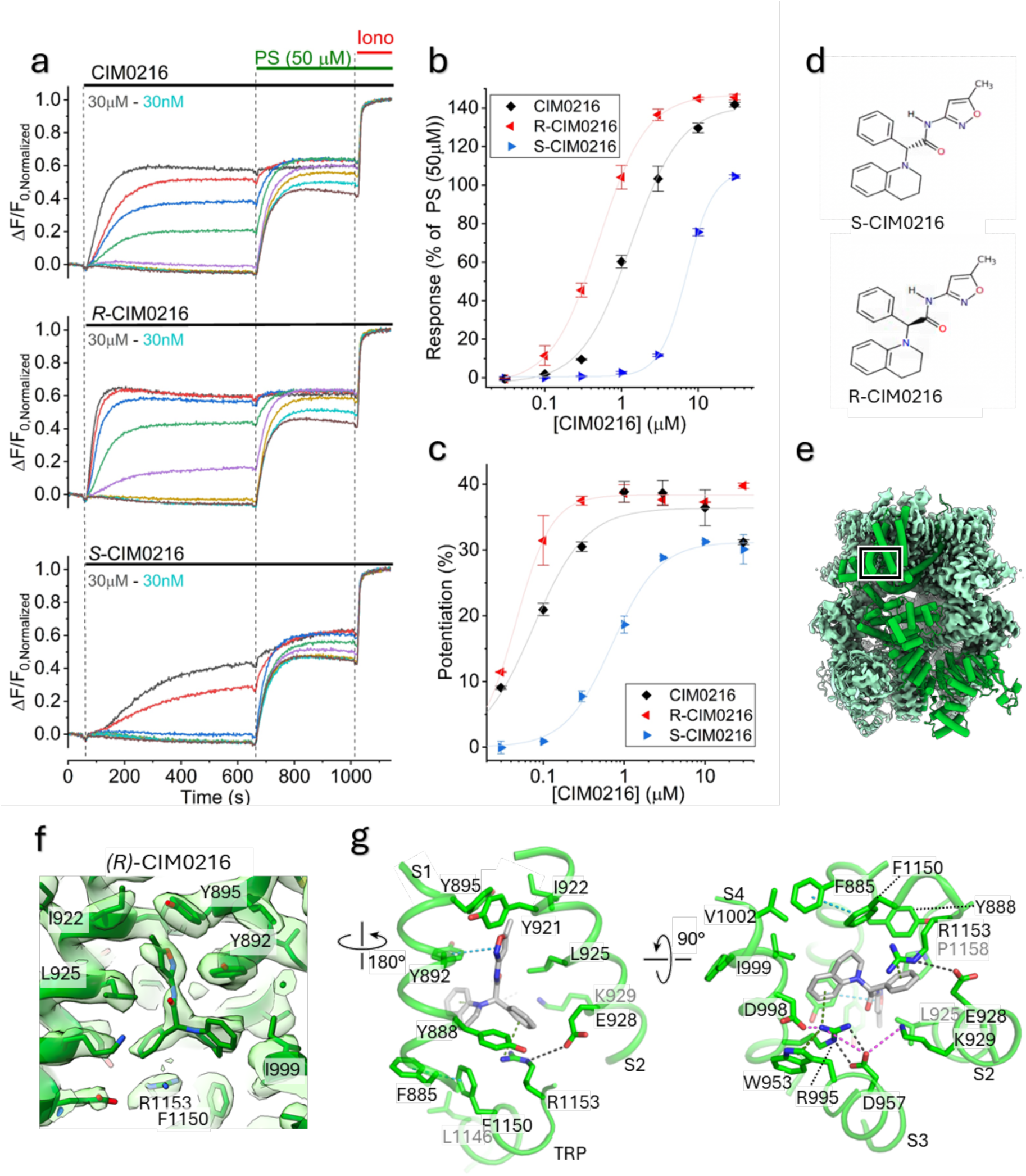
Stereoselectivity and structural basis of CIM0216 agonistic action. a) Normalized fluorescence traces showing changes in jRCAMP1b fluorescence in cells expressing WT TRPM3 upon stimulation with various concentrations of racemic CIM0216 and its two enantiomers followed by a stimulation by PS (50 μM). At the end of the experiment, a saturating ionomycin response was evoked for normalisation. b) Concentration-dependent activation curves showing the response to racemic CIM0216 and its two enantiomers, normalized to the maximal response at the end of the PS-stimulus. Full lines represent fits of the modified Hill function. c) Concentration-dependent potentiation by racemic CIM0216 and its two enantiomers. Potentiation was determined from the normalized fluorescence at the end of the PS-stimulus in the presence of racemic CIM0216 and its two enantiomers compared normalized to vehicle. d) Chemical structure of *(R)-* and *(S)-*CIM0216. e) Surface representation of TRPM3 in green in the presence of R-CIM0216, PS and PI(4,5)P2 at ambient temperature, with one subunit shown as cylinders. The CIM0216 binding site is highlighted with a box. Contouration in ChimeraX is at 0.053 σ. f) Close-up view on the binding site of R-CIM0216 together with the cryo-EM density. The protein is displayed in cartoon representation with residues shown as sticks. Contouration of the map in ChimeraX is at 0.165 σ g) Residues that coordinate R-CIM0216 are shown as sticks and labelled. Interactions with R-CIM0216 are shown as dashed lines; hydrogen bonds in black; salt bridges in pink; cation-π stacks in green and π-π stacks in cyan. Hydrophobic contacts are not specifically marked.

To obtain improved structural insights into the binding of agonists and events that lead to channel opening, we performed cryo-EM single-particle analysis of mouse TRPM3 samples in the combined presence of pure *(R)-*CIM0216, PS and the lipid phosphatidylinositol-(4,5)-biphosphate (PI(4,5)P_2_) at ambient temperatures. The final 3D reconstruction of TRPM3 in presence of agonists (TRPM3_AGO_) was resolved to 3.07 Å, and allowed us to identify clear EM densities corresponding to *(R)-*CIM0216 and PS (Figure 3E,F, Supplementary Figure 2-4, Supplementary Table 1). Whereas the binding pose for PS matches well with the site 1 in the recently described structure ^25^ (Supplementary Figure 7E and 10A-C), our results point at a very different orientation of *(R)-*CIM0216 in the binding pocket. In particular, we find that the methylisoxazole moiety of *(R)-*CIM0216 points upwards towards the extracellular side, making a π-π stack with residue Y892 (S1) and multiple hydrophobic contacts with the side chains of residues Y895 of S1 and Y921, I922 and L925 of S2, respectively; the phenyl moiety points sideways, and is sandwiched between residues in S1 (Y888), S2 (L925, E928) and the TRP domain (R1153). R1153 of the TRP domain builds a cation-π stack with the phenol group of CIM0216 and is positioned through E928; the tetrahydroquinalone moiety makes a cation-π stack with R995 (S4) and hydrophobic interactions with multiple residues in S1 (Y892) and S4 (R995, D998, I999) and in the post-S6 helix TRP domain (F1150). R995 is positioned through a network of interactions involving cation-π interactions with W953 and salt bridges as well as hydrogen bonds with D957, K929 and D998 (Figure 3G, Supplementary Figure 7D). TRPM3_AGO_ adopts a very similar fold as TRPM3_APO_ characterized by minor conformational changes with r.m.s.d values of less than 0.6 Å in average, suggesting a non-conducting conformation (Supplementary Figure 5C,D and 10E-G).

### Functional characterisation of the common ligand binding pocket

From the available cryo-EM structures of TRPM3, it is apparent that the antagonists *(R)-*isosakuranetin, ononetin and primidone, and the synthetic agonist and potentiator *(R)-*CIM0216 show partly overlapping binding poses with the channel, in a common binding pocket located between transmembrane domains S1-S4 and the TRP domain (Figure 1 and 3). Compared to the central position of primidone, both *(R)-*Isosakuranetin and Ononetin are extending further towards the cytosolic side, whereas the methylisoxazole moiety of *(R)-*CIM0216 points further upwards towards the extracellular side (Supplementary Figure 8).

To obtain a better understanding of the importance of specific ligand-amino acid interactions in this cavity on channel activity, we mutated residues surrounding the common binding pocket, and tested the impact of these mutants on the concentration-dependent effects of the different ligands in the jRCAMP1b-based calcium imaging assay. For these functional experiments we used the human TRPM3 consensus clone (NM_001366145.2), allowing direct juxtaposition of the structural data with human disease-associated variants ^11^ (Supplementary Figure 11; Supplementary Table 2). Residues were initially mutated to alanines; in cases where this led to a poorly functional channel we also introduced conservative mutations (e.g. K929R and R1153K). In total, we probed the functional effects of mutagenesis at 18 unique residues in the ligand-binding pocket, and found that sizable agonist responses could be measured for mutations at 13 residues (Supplementary Figure 12). For the mutants, calcium signals were measured during a 10-minutes exposure to a range of concentrations of the different ligands (*(R)-*CIM0216, primidone, isosakuranetin, ononetin and PS), followed by an invariant test stimulation with PS (50 μM) and a final ionomycin control stimulus for normalisation (Figure 4; Supplementary Figure 9). Note that for these experiments, we initially used commercial isosakuranetin, which in our analyses contained ∼50% of the active *(R)-*enantiomer (Figure 2). Where indicated, specific pure isosakuranetin enantiomers were used.

**Figure 4.**
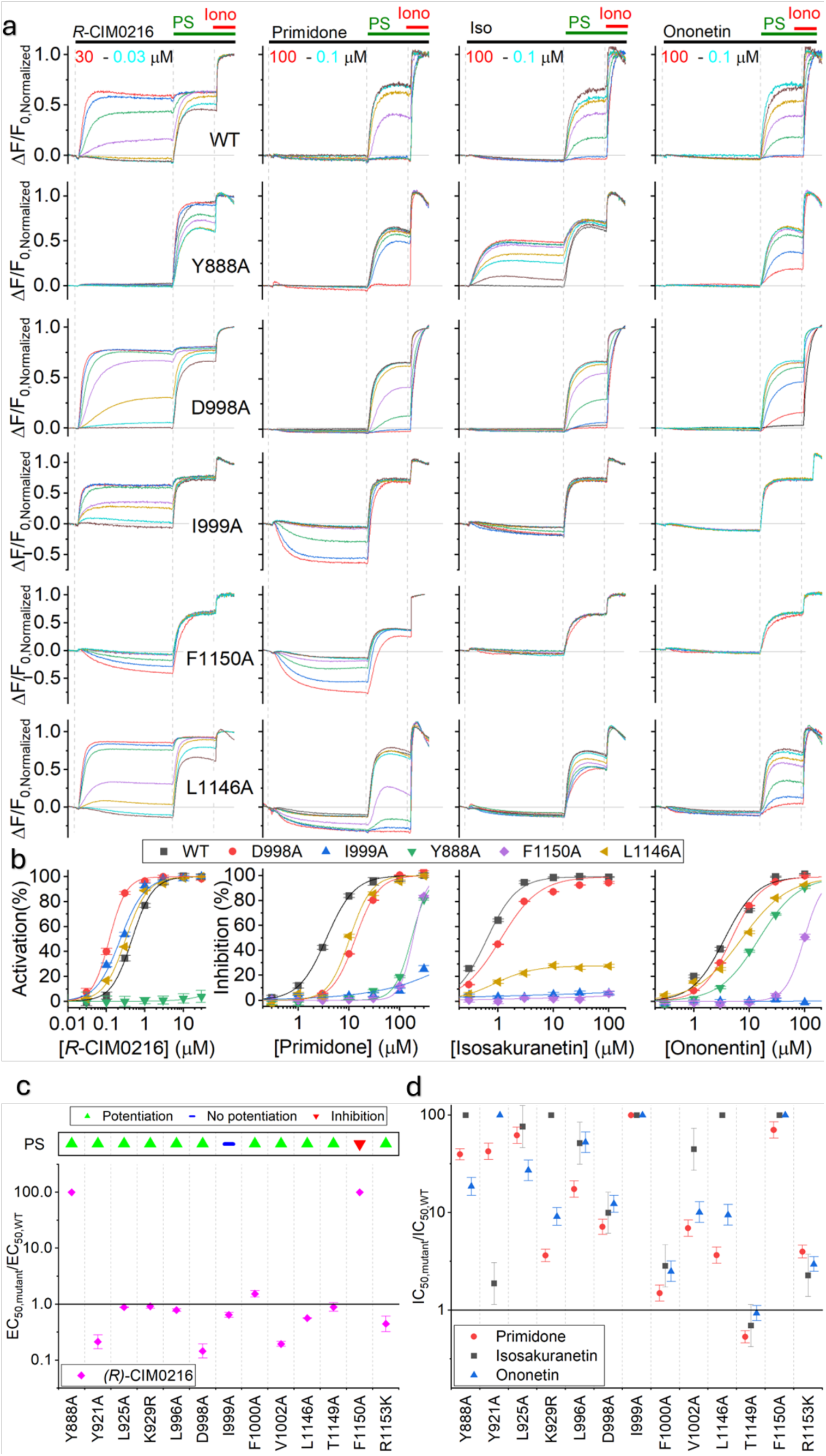
Mutations in the ligand-binding pocket differentially affect the effects of ligands. a) *(top)* Normalized fluorescence traces showing changes in jRCAMP1b fluorescence in cells expressing WT TRPM3 and the indicated mutants upon addition of various concentrations of the indicated ligand (*(R)*-CIM0216, primidone, isosakuranetin and ononetin) followed by a stimulation with PS (50 μM). At the end of the experiment, a saturating ionomycin response was evoked for normalisation. (bottom) Concentration-response curves for activation by *(R)*-CIM0216 or inhibition by primidone, isosakuranetin and ononetin for the indicated mutants. b) Summary of the effects of the indicated mutants on the inhibitory effect of the antagonists primidone, isosakuranetin and ononetin. Indicated values are the IC_50_ values for the respective compounds and mutants normalized to the IC_50_ values obtained for WT TRPM3. c) Same as (b), but for the effect of mutants on the EC_50_ values for the agonistic effect of *(R)-* CIM0216.

For wild type (WT) TRPM3, PS evoked concentration-dependent responses with an EC_50_ value of 5.8 ± 2.5 μM, in line with earlier studies^1,2^ (Supplementary Figure 13). In agreement with the notion that the PS binding site (Site 1) is located on the extracellular side of the channel, away from the ligand binding pocket, we found that most mutants showed relatively normal responses to PS, with EC_50_ values ranging between ∼1 and 15 μM (Supplementary Figure 13). In WT, as outlined above, *(R)-*CIM0216 activated WT TRPM3 and potentiated the PS response with EC_50_ values of ∼500 and ∼50 nM (see Figure 3), whereas isosakuranetin caused a concentration-dependent inhibition of the test response to PS (50 μM). Similarly, primidone and ononetin inhibited the PS response with IC_50_ values of 3 and 1 μM, respectively, in line with earlier studies ^20–22^.

In accordance with the structural insight for the interactions with each of the ligands described above, mutations at most residues flanking the ligand binding pocket caused marked and distinct alterations of the ligand response profile, examples of which as shown in Figure 4A. First, two mutations (Y888A and F1150A) largely eliminated the agonistic effects of *(R)-*CIM0216, showing no detectable increase in Ca^2+^ at the highest tested concentration (30 μM). Notably, for Y888A *(R)-*CIM0216 still caused a concentration-dependent potentiation of the PS response, with an EC_50_ value of 1.2 μM, whereas for F1150A, we measured neither activation nor potentiation, but observed a concentration-dependent decrease of the basal fluorescence. These findings indicate that these two residues, which interact with the tetrahydroquinalone (F1150 and Y888) and phenyl (Y888) moieties of *(R)-*CIM0216, are not absolutely essential for the binding of the ligand to the pocket, but crucially determine the effect downstream of ligand binding. Oppositely, several mutations (Y921A, D998A) caused a significant, >5-fold reduction in EC_50_ for *(R)-*CIM0216, indicative of an increased sensitivity for the ligand. Notably, the side chain of Y921 points towards the ligand binding pocket in the APO and antagonist-bound structures, but when *(R)-*CIM0216 is bound it rotates outwardly, avoiding a clash with the methylisoxazole moiety of the ligand (Supplementary Figure 14). Such outward rotation would not be necessary to accommodate *(R)-*CIM0216 in the Y921A mutant. In addition, several mutations significantly reduced the inhibitory potency of the tested antagonists. These included mutations that largely abolished the inhibitory effect of all three antagonists, such as I999A, F1150A, corroborating the interactions between these residues and all three antagonists in the cryo-EM structures in the central part of the binding pocket. Other mutations had a more specific effect on antagonist action. K929R and L1146A largely abolished the inhibitory effect isosakuranetin, but only mildly affected inhibition by primidone. These effects are in line with the cryo-EM structures showing that isosakuranetin is positioned lower (i.e. closer to the cytosolic side) in the binding pocket, where it can interact with K929 and L1146, whereas the smaller primidone molecule does not reach far enough towards the cytosolic side to interact with L1146 or form equally strong bonding with K929. Oppositely, Y921A strongly reduced the inhibitory potency of primidone, whereas inhibition by isosakuranetin was not significantly affected. In the cryo-EM structures, Y921 is situated in the upper part of the binding pocket, where it forms a hydrogen bond with the pyrimidinedione ring of primidone with its phenol ring, while not directly interacting with the lower positioned isosakuranetin. The impact of these mutations on the inhibitory effects of ononetin were mostly intermediate to those on isosakuranetin and primidone. Figure 4B,C summarizes the effects of the different mutations on ligand potency, as the fold-change in EC_50_ values for activation by *(R)-*CIM0216 or in IC_50_ values for the antagonists, as compared to the EC_50_/IC_50_ values for WT. Taken together, the results from the mutagenesis studies corroborate the ligand binding modes for antagonists and *(R)-*CIM0216 determined using cryo-EM.

### Specific ligand-channel interactions dictate the functional outcome of ligand action

When analysing the results obtained with mutations Y888A and F1150A (Figure 4A), we observed unexpected effects for some of the tested ligands. In the case of Y888A, we found that isosakuranetin, instead of antagonizing responses to PS, acted as a robust, concentration-dependent agonist, with an EC_50_ value of 3 μM. Notably, when we tested the two isosakuranetin enantiomers separately, we found that *(S)-*isosakuranetin, which is inactive at the wild type channel, acts as a potent agonist of the Y888A mutant (EC_50_ ∼600 nM), while *(R)-*isosakuranetin partially inhibits the mutant channel, albeit with a lower potency than for the wild type channel (Figure 5). Interestingly, in the APO structure we observed two rotamers for Y888, with one where the tyrosine side chain points towards the center of the ligand binding pocket, and the other where it is rotated away from the center (Supplementary Figure 15). In the structures where the ligand binding pocket is occupied by *(R)-*CIM0216, primidone, isosakuranetin or ononetin, only the latter rotamer is observed, preventing a clash between the tyrosine side chain and the ligands. To further explore the importance of Y888 for channel gating and ligand responses, we tested the effect of additional aromatic and non-aromatic substitutions at this position. Substituting tyrosine by the polar, non-aromatic amino acid serine (Y888S) had mostly similar effects as seen for Y888A: the mutation fully abrogates the inhibitory effect of primidone and the agonistic effect of *(R)-* CIM0216, preserves *(R)-*CIM0216-induced potentiation of the PS response and reduces the inhibitory potency of ononetin. With regards to isosakuranetin, the *(S)*-enantiomer acted as an agonist and *(R)-* isosakuranetin as an antagonist for Y888S. No agonistic effects were seen for *(S)*-isosakuranetin upon mutation of Y888 to the aromatic residues histidine or phenylalanine. The polar aromatic amino acid histidine (Y888H) conserved activation and potentiation by *(R)-*CIM0216 and inhibition by *(R)-* isosakuranetin and ononetin, albeit with reduced potency, but abolishes the antagonistic effects of primidone. The non-polar aromatic amino acid phenylalanine (Y888F), largely conserved the inhibitory effects of *(R)-*isosakuranetin and primidone, as well as the agonist and potentiating effects of *(R)-* CIM0216 its agonist effects (albeit with higher IC_50_/EC_50_ values).

**Figure 5.**
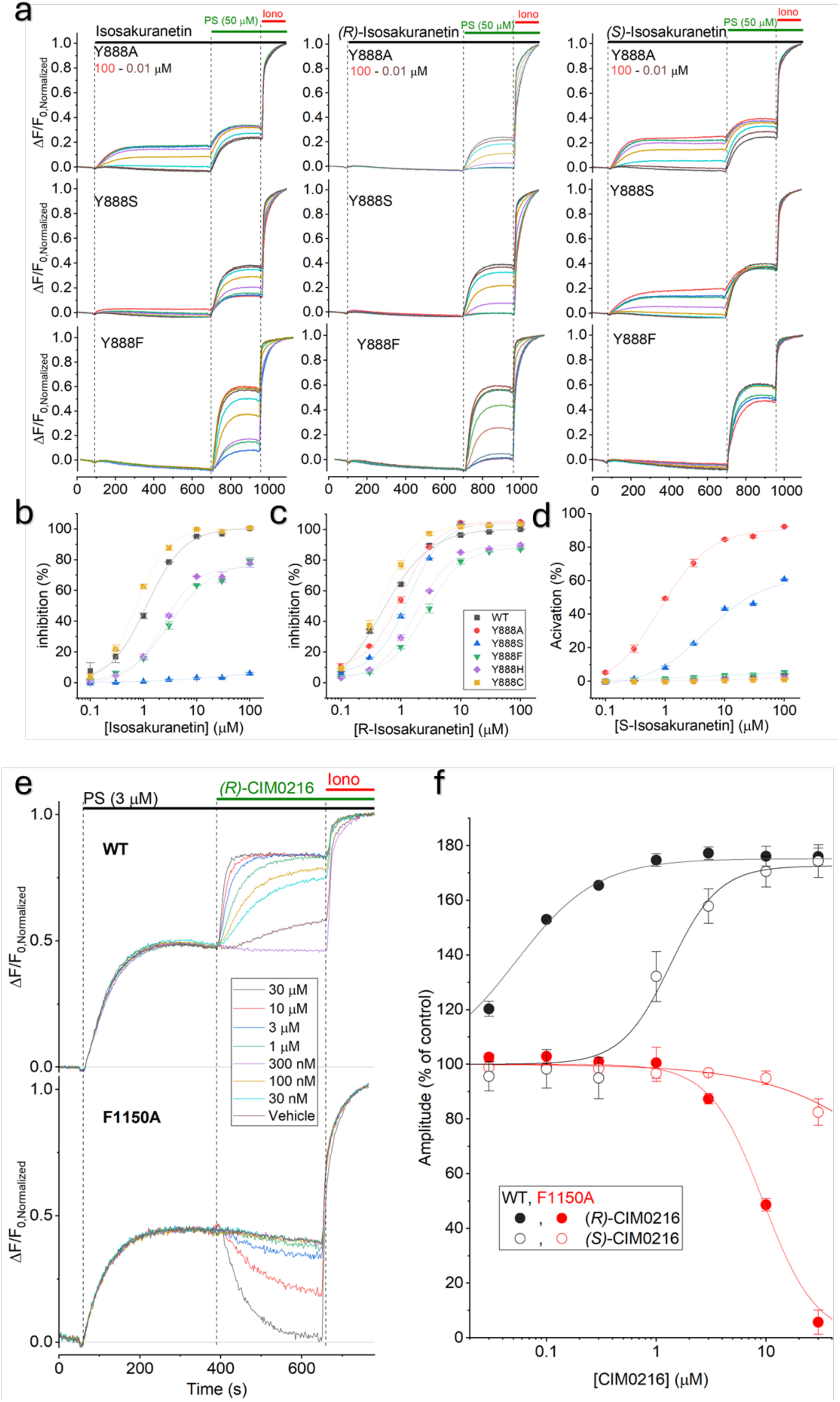
Mutating residues Y888 and F1150 affects the directionality of ligand action. a) Representative jRCAMP1b fluorescence assays showing the effect of racemic isosakuranetin and the *(R)*- and *(S)*-enantiomers on mutations at residue Y888. b) Concentration dependence of the inhibitory effect of racemic isosakuranetin at WT and the indicated Y888 mutants. Note that Y888A was not inhibited. c) Concentration dependence of the inhibitory effect of *(S)*-isosakuranetin, showing inhibition of all tested Y888 mutans. d) Concentration dependence of agonistic effect of *(R)*-isosakuranetin, which acts as an agonist at Y888A and Y888S, but not at other Y888 mutants. e) Representative jRCAMP1b fluorescence assays showing the concentration-dependent effects of (R)-CIM0216 on the response of WT and F1150A to a submaximal concentration of PS (3 μM). f) Summary of the concentration dependence of the effects of CIM0216 enantiomers on the response of WT and F1150A to PS (3 μM), showing potentiation for the former and inhibition for the latter.

In the case of F1150A, we found that *(R)-*CIM0216, instead of evoking a calcium response, caused a concentration-dependent reduction in the basal jRCaMP1b fluorescence, indicative of inhibition of the channel’s basal activity (Figure 4). Moreover, at the highest concentrations (10 and 30 μM), we also observed a partial inhibition of the subsequent response to 50 μM PS (Figure 4). To investigate this inhibitory effect further, we performed experiments where cells were first stimulated with a sub-maximal concentration of PS (3 μM), and subsequently exposed to *(R)-*CIM0216 at concentrations between 30 nM and 30 μM. For WT, PS evoked a robust response, followed by a concentration-dependent further activation by *(R)-*CIM0216, with an EC_50_ of 53 ± 5 nM (Figure 5), similar to the potentiating effect described in Figure 3. In contrast, for F1150A, *(R)-*CIM0216 caused a concentration-dependent inhibition of the PS-evoked calcium signal, with an IC_50_ value of 9.4 ± 6 μM (Figure 5). Notably, *(S)*-CIM0216 did not show any detectable activatory or inhibitory effect on F1150A.

These findings reveal that subtle changes in the binding pocket and the stereochemistry of the ligands not only affect the ligands’ potency but also change the directionality of their effects, turning an agonist into an antagonist or *vice versa*.

### Novel patient mutations affecting the ligand binding pocket

Gain-of-function variants in the TRPM3 gene underly a spectrum of neurodevelopmental disorders, with developmental delay/intellectual disability and epilepsy as dominant phenotypic features ^17–19^. Since increased TRPM3 channel activity likely underlies the neurological symptoms, experimental treatment with primidone has been initiated for several patients, generally leading to a noticeable improvement in development and electrophysiological parameters ^19,23^. We identified two individuals with epilepsy and EEG anomalies, who were found to be heterozygous for two different, previously undescribed *de novo* variants in the TRPM3 gene, namely I999S and F1150S. Figure 6A,B provides an overview on the location of these and previously identified gain-of-function patient mutations. A clinical summary of the two new patients can be found in Supplementary Table 3. While the treating physicians considered initiating treatment with primidone, we realized that the specific disease-associated variants affect residues in the ligand binding pocket, at sites where alanine mutations (I999A and F1150A) had an important impact on ligand effects. We therefore evaluated the activity and ligand responses of both clinical variants.

**Figure 6:**
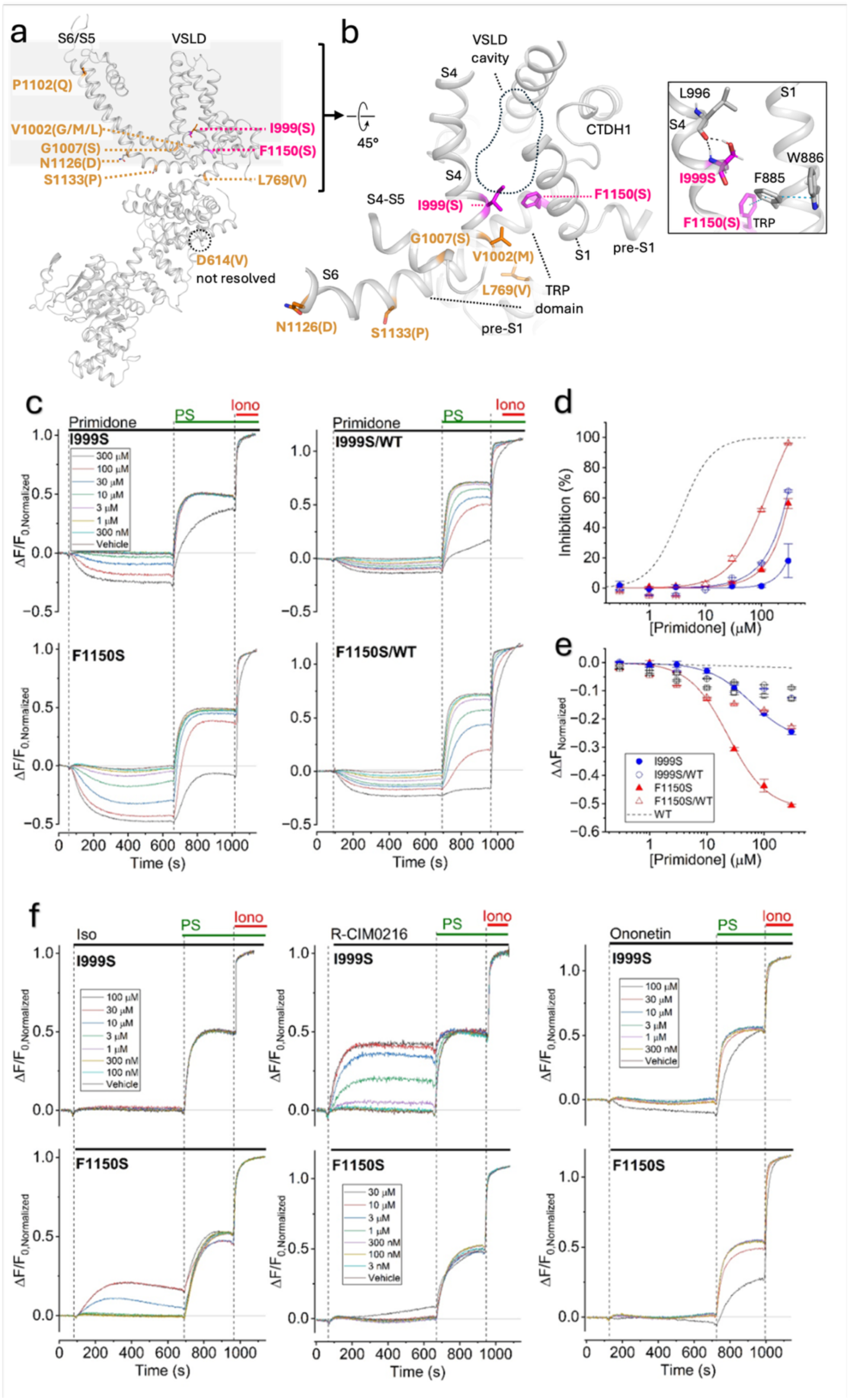
D*e novo* mutations in patients with epilepsy affecting the ligand-binding pocket. a) Plot of known (orange) and new (pink) disease-causing variants in patients with epilepsy and neurodevelopmental disoreders on the structure. Only one subunit is shown in cartoon mode and side chains for the residues at the gain-of-function mutant position are shown as sticks. Gain-of-function mutants refer to the numbering in the human TRPM3a2 isoform. b) Close-up view of a). The new mutations, I999S and F1150S, contribute to the ligand-binding pocket. Close-up view on I999 and F1150 (right). A π-π stack between F1150 and F885 would be abrogated in the F1150S mutant. The I999S mutation could form a hydrogen bond between its serine side chain hydroxyl group and the main chain carbonyl oxygen of L996, potentially affecting the stability of the S4 helix by inducing a kink. c) Representative jRCAMP1b fluorescence assays showing the effect of primidone in cells expressing I999S and F1150S, either alone or in a 1-to-1 ratio with WT TRPM3. d) Concentration dependence of the inhibitory effect of primidone on the PS response for the conditions shown in (c). The dotted line indicates the concentration-response curve for WT TRPM3. e) Concentration dependence of the effect of primidone on the basal fluorescence for the conditions shown in (c). f) Representative traces showing the effects of other ligands (isosakuranetin, (R)-CIM0216 and ononetin) on I999S and F1150S. Note the severely affected inhibition by isosakuranetin and ononetin, the insensitivity of F1150S for CIM0216, and the activaition of F1150S by isosakuranetin.

Similar to other disease-associated TRPM3 variants, I999S and F1150S exhibited gain-of-function characteristics, reflected in higher basal jRCAMP1b fluorescence levels compared to WT and lower EC_50_ values for PS activation (Supplementary Figure 13 & 16). Importantly, I999S and F1150S severely affected the channel’s sensitivity to primidone, causing a >100-fold increase in IC_50_, and less than 50% inhibition of the PS-induced response at the highest tested concentration (300 μM; Figure 6). To mimic the condition of the patients, which are heterozygous for the mutations and thus likely express comparable levels of wild type and mutant subunits, we performed experiments where we co-expressed WT and mutant TRPM3 in a 1-to-1 ratio. Under this condition, we still observed a robust reduction in sensitivity to primidone inhibition, with IC_50_ values of 85.2 ± 9.1 μM and 222 ± 15 μM for I999S/WT and F1150S/WT, respectively (Figure 6). As such, treatment of patients carrying these mutations with primidone, which is currently the only available targeted therapy for patients with TRPM3-related neurodevelopmental disorders, may not be as effective as in patients that have gain-of-function variants outside the ligand binding pocket. Interestingly, primidone also caused a concentration-dependent reduction of the non-stimulated jRCAMP1b fluorescence for both I999S and F1150S, indicating inhibition of basal channel activity. Estimated IC_50_ values for the reduction of basal activity were 66.1 ± 3.6 μM and 23.4 ± 4.9 μM for I999S and F1150S, and 5.4 ± 1.1 μM and 3.5 ± 0.3 μM for I999S/WT and F1150S/WT respectively.

We further tested the effects of the other ligands on the two disease mutants. F1150S was insensitive to *(R)-*CIM0216 (i.e. no detectable activation, potentiation or inhibition), which contrasts to I999S and other disease-associated mutations (e.g. V1002M, P1102Q) that exhibit robust activation by CIM0216 ^15^ (Figure 6). In addition, both I999S and F1150S showed a strongly reduced sensitivity to inhibition by ononetin and isosakuranetin (Figure 6). Notably, for F1150S we even observed activation by isosakuranetin at concentrations ≥10 μM (Figure 6). Taken together, these findings indicate that disease-associated variants affecting the ligand binding pocket have a strong impact on the effect of ligands.

## Discussion

In this study, we provide important new insights into a ligand-binding pocket in TRPM3, located in the VSLD and lined by residues from S1-S4 and the C-terminal TRP box. Based on high-resolution cryo-EM structures of TRPM3 in the apo state and in complex with different antagonists (primidone, *(R)-* isosakuranetin, ononetin) and with the agonist and potentiator *(R)-*CIM0216, we obtained a detailed picture of the interaction mode of these ligands within the ligand-binding specific residues in the channel. We found primidone to be located at the centre of the VLSD binding pocket. Compared with a recent study ^25^, primidone is tilted by 45° around the axis of the phenyl group and coordinated through an extensive network of residues that form additional and different contacts. Hence, the binding pose deviates significantly compared to the previously published data ^25^ (Figure 1 and Supplementary Figure 6). The observed discrepancies regarding the primidone binding site could be attributed to a less defined and therefore ambiguous density in the primidone binding pocket of 9B28 despite very similar reported resolutions (Supplementary Figure 6B).

Primidone has been evaluated in small scale clinical studies in the treatment of TRPM3 gain-of-function mutants where it could attenuate electroencephalographic abnormalities. Hence, this compound could serve as a point of departure for the rational design of more potent drugs with better controlled side-effects. A thorough understanding of the precise interaction in the ligand-binding pocket is thus critical for such progress in particular because of the promiscuous binding pocket of TRPM3 where minimal structural differences between ligands may result in agonistic or antagonistic effects. In comparison, the binding poses of *(R)-*isosakuranetin and ononetin show overlap with that of primidone, but these larger molecules, which exhibit a higher apparent affinity for the channel, make additional interactions extending further towards the cytosolic side, namely with E928 and E932 (S2), V1002 (S4) and L1146 (TRP domain). *(R)-*CIM0216 also occupies the centre of the binding pocket, with its methylisoxazole moiety pointing much further upwards towards the extracellular side compared to the antagonists (Supplementary Figure 8). We performed a comprehensive functional analysis of point mutations in the ligand binding pocket on the potency of the different ligands (IC_50_/EC_50_), confirming the significance and specificity of different residues for ligand-induced channel modulation.

A key insight from our study is the highly stereoselective action of isosakuranetin and CIM0216 on TRPM3. Previous publications and curated online pharmacology resources attributed the antagonistic effect of isosakuranetin on TRPM3 to the *(S)*-enantiomer ^20,31^, which is the expected enantiomer produced in the highly stereoselective flavanoid biosynthetic pathway in plants ^29,30^. In contrast, our structural and functional data demonstrate that it is in fact *(R)*-isosakuranetin that potently inhibits TRPM3, while the *(S)*-enantiomer is largely inactive on the wild-type channel. Intriguingly, the *(S)*-enantiomer acts as a potent agonist of the Y888A and Y888S mutants. Our findings indicate that commercially available, plant-derived isosakuranetin from different sources contains a racemic mixture of *(R)*- and *(S)*-isosakuranetin. This may seem at odd with the stereoselectivity of the biosynthetic pathway in plants producing primarily (2S) flavanones. Possibly, epimerization of *(S)*-isosakuranetin may occur after its production in the plants or during the purification process. These findings also imply that most -if not all-previous *in vitro* and *in vivo* studies that involved isosakuranetin as a TRPM3 antagonists used a mixture of *(R)-* and *(S)-*isosakuranetin. Notably, in studies on the role of TRPM3 in pathological pain, both TRPM3-dependent and TRPM3-independent analgesic effects of isosakuranetin have been reported. Even if the TRPM3-independent molecular targets of isosakuranetin and their stereoselectivity are currently unknown, the use of pure *(R)*-isosakuranetin may be advantageous in future studies to obtain higher efficacy at lower doses, possibly with reduced side-effects.

Our results also reveal the stereoselectivity and binding mode of the potent synthetic agonist and potentiator CIM0216. Whereas this compound, which contains one chiral center, has previously only been used as a racemic mixture, we performed chiral HPLC and vibrational circular dichroism analysis to separate the pure (*R)-* and (*S)-*enantiomers of CIM0216. We found that the agonistic and potentiator effects of racemic CIM0216 are primarily driven by *(R)-*CIM0216, and that the potency of *(S)-*CIM0216 is >20-fold lower than that of *(R)-*CIM0216. Moreover, by using the pure (*R)-*enantiomer during the cryo-EM structural determination, we were able to obtain a high-resolution EM density for *(R)-* CIM0216, which allowed us to model the binding pose of the correct compound in the VSLD with high fidelity. This binding pose differs substantially from a previously proposed model, which was based on a low-resolution cryo-EM structure (obtained using racemic CIM0216) and all-atom molecular dynamics simulations that assumed the inactive (*S)-*enantiomer ^25^. In contrast to that model, we find that *(R)-*CIM0216 is positioned higher up in the VSLD, with the methylisoxazole moiety pointing towards the extracellular side. More specifically, the methylisoxazole moiety of *(R)-*CIM0216 fills the space that is occupied by the phenol side chain of tyrosine 921 in the APO and antagonists-bound structures, leading to an up- and sidewards rotation of the side chain of tyrosine 921. We speculate that this motion may contribute to channel activation and/or potentiation induced by *(R)-*CIM0216 binding. Notably, substituting an alanine for tyrosine 921 (mutant Y921A) shifted the concentration-activation curve for *(R)-*CIM0216 towards lower concentrations, further highlighting the importance of this residue for channel *(R)-*CIM0216 activity.

An overall conclusion of our study is that the ligand binding pocket in the VSLD of TRPM3 exhibits a high degree of ligand promiscuity. Indeed, the same cavity forms the interaction side for both activating and inhibitory ligands, and apparently subtle changes to ligand or binding pocket can change the directionality of the effect of ligand binding on channel gating. This is exemplified by the mutation F1150A, which turned *(R)-*CIM0216 into an antagonist, reducing basal channel activity and PS-evoked responses, and by mutations Y888A, Y888S and F1150S, which caused *(S)*-isosakuranetin to act as a concentration-dependent agonist. These findings indicate that mutations at these residues not only influence binding of the ligands to their binding site, but also affect the conformational changes that couple ligand binding in the VSLD to channel gating. The promiscuity and stereoselectivity of the ligand binding site adds a layer of complexity to structure-aided design approaches to develop novel pharmacological tools targeting TRPM3. Indeed, based on available data it is difficult to predict whether compounds that bind to the ligand binding pocket will promote or antagonize channel opening, and subtle changes in ligand or channel can potentially affect the directionality of the effect. This duality of the binding pocket requires further studies to be entirely understood, such as a structure of the fully conductive state, but it is already clear that this genuine feature has to be taken into account for the rational design of novel drugs and for the treatment of pathological gain-of-function mutant carriers.

In recent years it has been established that rare gain-of-function variants of TRPM3 underlie a spectrum of neurodevelopmental disorders in human. At this point, about a dozen different mutations - mostly *de novo -* have been reported in patients, invariably leading to a dominant gain of channel function phenotype when expressed in heterologous expression systems. Primidone, which inhibits wild type TRPM3 with an IC_50_ value of ∼3 μM in our assay (values in the literature range between 0.6 and 5 μM)^14,15,22,23^, can provoke substantial inhibition of TRPM3 function at typical therapeutic plasma levels (∼20 μM) measured in patients using this clinically approved drug for indications such as epilepsy or essential tremor^32^. Currently, several patients with neurodevelopmental disorders due to TRPM3 gain of function mutations are being treated off-label with primidone, with highly encouraging improvements regarding neurodevelopment and seizure control^19,23^. In this study, we described two new missense TRPM3 variants in two patients with epilepsy and neurodevelopmental delay, introducing amino acid substitutions at critical residues in the primidone binding site, namely I999S and F1150S. The substitution of I999 by a serine could create polar interactions with close-by residues or cause a kink in the S4-helix, while the F1150S mutation on the other hand would abrogate a π-π stacking network that we observe between F1150, F885 and W886 presumably leading to a destabilization in this region (Figure 6B, right). These amino acid substitutions could promote a channel configuration that has a higher propensity to open. Importantly, these disease mutations not only cause a gain of channel function, but also severely affect the antagonist action of primidone, with IC_50_ values ≥100 μM for the heterozygous (I999S/WT and F1150S/WT) conditions. These findings need to be taken into account when planning pharmacotherapy for these patients, as primidone levels obtained with regular treatment regiments are unlikely to have any clinical effect, in contrast to the beneficial effects found in patients carrying TRPM3 gain-of-function variants affecting other regions of the channel.

Like in previous structural work on TRPM3 ^24,25^, the obtained cryo-EM structures all show the pore in the closed state, even when the structure was obtained in the combined presence of PS, PI(4,5)P_2_ as well as the potent agonist and potentiator *(R)*-CIM0216, and at room temperature. Therefore, our structures do not reveal the specific conformational changes associated with channel opening/closing following ligand binding in the VSLD domain. Interestingly, the antagonist-bound structures reveal specific polar interactions between residues lining the VSLD with the probed antagonists while for the agonist and potentiator *(R)*-CIM0216, we find largely non-polar side chains that mediate hydrophobic contacts (Supplementary Figure 7). Based on the type of interactions, *(R)*-CIM0216 could act as a molecular lubricant in the binding pocket facilitating movements towards channel opening, while the antagonists that make numerous, strong polar interactions inside the VSLD could function as a molecular glue to prevent the conformational changes required during channel activation. Possibly, the absence of a supporting biomembrane and of a specific transmembrane potential (TRPM3 is activated upon depolarization to potentials >+0mV) may result in an abundance of particles containing a non-conducting conformation of the channel.

## Acknowledgements

We thank Drs. Marcus Fislage and Dirk Reiter (Biogenic Electron Cryo-Microscopy (BECM), Brussels) for support with cryo-EM data collection and technical assistance with workstations, and Sara Kerselaers and Melissa Benoit for technical assistance. Research infrastructure was funded by the KU Leuven Research Council (AKUL/19/34). This work was supported by grants from the Research Foundation — Flanders (FWO; G0B9520N, G0B7620N and G055124N to T.V.), the KU Leuven (C24M/21/028 to T.V.), the Queen Elisabeth Medical Foundation for Neurosciences (to T.V.) and the Flemish Institute for Biotechnology (VIB to T.V. and J.D.B.).

## Contributions

TV and JDB conceived the study and wrote the manuscript. BB, AVS, SS, TV and JDB planned experiments; BB performed and analyzed all the functional experiments; SS cloned constructs used for cryo-EM and generated the stable cell line expressing murine TRPM3. SS and JDB expressed, purified and prepared proteins for cryo-EM. AVS prepared graphene grids, collected cryo-EM data and made initial models. AVS and JDB processed cryo-EM data. JDB built atomic models and deposited them. BB, SS, AVS, TV and JDB analyzed the data. RR, JV and TV procured patient data. AJ and BB designed constructs for expression of wild type and mutant human TRPM3. JV and TV performed pilot experiments on the activity of CIM0216 enantiomers.

## Materials and methods

### TRPM3 expression and purification for structural work

The coding sequence of murine TRPM3 (NCBI Reference Sequence: NP_001030319.1), truncated at amino acid residue 1344, was cloned into a modified pcDNA5/FRT/TR expression vector (Invitrogen/Thermo Scientific). To do so, the entire expression cassette of pcDXC3GMS (Addgene #49031) containing a C-terminal SBP-tag was excised with HindIII and ApaI and ligated into HindIII/ApaI-digested pcDNA5/FRT/TR to generate p5TO-CSBP. Thereafter, SapI-flanked PCR products of TRPM3 were ligated into p5TO-CSBP by FX cloning ^33^. Tetracycline-inducible stable cell lines were obtained following the manufacturer’s instructions (Flp-In T-REx293 cell manual, Invitrogen/Thermo Scientific) and as described earlier ^34^,after a co-transfection of Flp-In T-REx293 cells with pOG44 and p5TO-CSBP-TRPM3 and following a selection at 100µg/ml hygromycin-B. Single colonies were picked, isolated and grown at 50µg/ml hygromycin-B for a minimum of four passages. Well expressing clones were identified by screening of purified TRPM3 on an Agilent 1260 HPLC by SEC with a Superose6 increase 5/150 column (Cytiva) and SDS-PAGE. Expression of TRPM3 was induced at a confluency of 60-80% with Dulbecco’s Modified Eagles Medium (DMEM, Sigma) containing 10% fetal bovine serum (TICO Europe) and 3µg/ml tetracycline. 24 hours after induction, 5-10 mM butyrate was added to boost protein expression. After a total of four days, the cells were harvested by centrifugation and frozen in liquid N2.

All subsequent purification steps were performed on ice. Frozen cell pellets were thawed and extracted with a buffer containing 50 mM Bicine pH 8.5, 200 mM NaCl, 10% glycerol, 1.5-2% glycodiosgenin (GDN) and protease inhibitors (cOmplete, Roche) for 1 hour. Aggregates and cell debris were removed by ultracentrifugation for 30 minutes at 130,000 g_av_ using a Beckman 45Ti rotor. The resulting supernatant containing TRPM3 was incubated with streptavidin resin (Thermo Scientific) for 1.5 hours. Unbound proteins were washed off using wash buffer (30 mM Hepes pH 7.5, 150 mM NaCl, 6% glycerol, 0.0063% GDN) and bound TRPM3 was eluted with wash buffer containing 10 mM biotin. Subsequently, the eluted protein was concentrated to 100µl using a 100 kDa cut-off concentrator (Amicon) and injected to a Superose6 increase 5/150 column (Cytiva) equilibrated with 10 mM Hepes pH 7.5, 150 mM NaCl and 0.0063% GDN. The peak fractions were collected and concentrated to 0.1-0.3 mg/ml.

### Preparation of cryo-EM grids

Graphene grids were prepared in-house using Trivial Transfer Graphene (ACS Material) or Graphene Easy Transfer (Graphenea) monolayer graphene sheet following a protocol introduced by Ahn et al ^35^. For this, Quantifoil R 2/1 Au 300 or 400 mesh or Quantifoil R 1.2/1.3 Au 300 mesh grids were utilized. Briefly, grids were transferred on a 3D-printed graphene transfer tool ^35^ immersed in the water in the Petri dish. The 2.5 cm x 2.5 cm PMMA/graphene pad was floated and carefully matched with the grids. The transfer tool was slowly lifted to coat the grids and dried in an oven preheated to 100 ℃ for one hour. Subsequently, individual grids were detached with a scalpel. PMMA was removed by immersing the grids in acetone (50 ℃, 30 min, repeated three times with mild stirring) followed by baking in the oven (preheated; 200 ℃, 6-7h). Graphene-coated PMMA-free graphene grids were stored in a desiccator and used within two months after their production. Grids were treated by glow discharging for 10-15 s at 0.3-0.35 mbar and with an applied current of 2.5-3 mA using an ELMO system (Cordouan technologies). 3 μl of TRPM3 in the apo-state or mixed with primidone (100µM) were loaded on the glow-discharged graphene grid, blotted immediately from the back side for 2 s at ambient temperatures and 90% relative humidity and flash frozen in liquid ethane using a Cryoplunge 3 System (Gatan). A volume of 2 or 3 μl of TRPM, mixed with 10 μM isosakuranetin or ononetin, or incubated with a combination of agonists (10μM CIM0216, 250 μM PregS, 100 μM PI(4,5)P_2_ extracted from porcine brain), was applied to the graphene grid and blotted immediately from the back side for 2.6-3.3 s. The chamber contained 90% relative humidity and was cooled to 6°C in the case of the antagonists or kept at 20°C in case of the agonist sample before flash freezing in liquid ethane using an EM GP2 plunger (Leica). All antagonists were incubated on ice with TRPM3 for about 20-40 min and plunged at 6°C, while the agonists were mixed with the protein for 30 min on ice before and incubation of 2 min at 30°C prior to plunging at 20°C. Grids were stored in liquid nitrogen before imaging.

### EM data acquisition

Images were collected at 300 kV on a CRYO ARM 300 (JEOL) electron microscope at a nominal magnification of 60,000 and corresponding pixel size of 0.69-0.71 Å. The images were recorded using a K3 detector (Gatan) operating in correlative-double sampling (CDS) mode. The energy filter slit was centered on the zero-loss peak with a slit width of 15 eV. 60 frames per movie were collected using SerialEM (SerialEM 4.1 beta 24 and 26, SerialEM 4.1.8, SerialEM 4.2 beta 8) ^36^ at an exposure time of 2.796 s, total dose of about 62 e-/Å2 and a defocus between 0.6-1.6 µm using a 3 × 3 or 5 x 3 multi-shot pattern for R1.2 or R2.1 grids, respectively ^37^. A total of 6,163, 11,614, 8,155, 7,812 and 18,427 movies were collected for TRPM3 in the apo state, with primidone, isosakuranetin, ononetin or with agonists (CIM0216, PregS, PI(4,5)P2), respectively.

### Image processing and 3D reconstruction

Data pre-processing was performed in cryoSPARC Live. Motion correction was done using Patch Motion correction and the contrast transfer function (CTF) of the resulting dose-weighted averages was estimated using CryoSPARC CTF estimation ^38^. All subsequent processing steps were performed in CryoSPARC v.4.5.3 ^38^. If required, manual curation was performed to exclude bad images. Picking was done with blob, Topaz and/or template picker using templates generated from published density of TRPM3 (EMD-28031) ^24^ or from selected class averages obtained after 2D classification. Initial processing was done using 4x binned particles (several rounds of 2D classification, *ab initio* modeling and hetero-refinement). Particles representing best classes were first binned twice, re-extracted and subjected to heterogenous, homogeneous and non-uniform refinement jobs, later particles were re-extracted without binning for the final reconstructions such as homogenous and non-uniform refinement jobs (see Supplementary Figure 1 and 2 for an accurate description).

### Model building and refinement

For model building of TRPM3_ONO_, the previously published cryo-EM structure of TRPM3 (PDB ID: 8DDS) ^24^ was rigid body fitted into the experimental cryo-EM map of TRPM3_ONO_ and refined in Coot v.0.9.8.2 ^39^. Using our cryo-EM density as a guide, regions that were not present in the previously published structure were either built *de novo* or taken from ColabFold ^40^ models of fragments, typically corresponding to rigid bodies in the structure. The resulting model was used as a custom template in ColabFold to relax amino acid side chain positions using AMBER molecular dynamics simulations. Stretches not visible in the density map were removed from the top ranked relaxed model. Further refinements were performed iteratively in Coot, ISOLDE ^41^ and Phenix (v.1.21-5207) ^42^ using a composite map of TRPM3_ONO_ obtained in Phenix from a locally refined map (N-terminus) and a consensus map from a homogenous refinement job.

The final model of TRPM3_ONO_ was used as a starting point to build the atomic models for TRPM3_ISO,_ TRPM3_APO,_ TRPM3_PRM,_ and TRPM3_AGO_ using the respective final maps (non-uniform and/or DeepEMhancer ^43^ maps). For this the model of TRPM3_ONO_ was first rigid-body fitted into the respective cryo-EM densities and side chains were adjusted accordingly. Residues not visible in the respective densities were removed, and residues were built *de novo* where the density allowed it. Refinements were done iteratively in Coot and Phenix until reaching good or excellent statistics. Initially, only TRPM3 monomers were refined while at later time points, monomers were assembled to tetramers for a last refinement step in Phenix. Restraints for ligands were generated from smiles codes using eLBOW from the Phenix suite or were retrieved in Coot from the monomer library in the case of phospholipids, cholesterol and detergent molecules. Compounds were initially fitted in the corresponding density using Coot and then refined in Phenix. In the case of TRPM3_AGO_ the PI(4,5)P2 molecule could not be modelled due to absence of an unambiguous density at the known PI(4,5)P2 binding site.

Figures to display atomic models and cryo-EM maps were prepared using Pymol 3.1.1 and ChimeraX-1.8. Interactions between TRPM3 and the ligands were analyzed using Pymol 3.1.1, the Protein-Ligand Interaction Profiler (PLIP) server ^44^ and LigPlot+^45^. The TRPM3 pore was analyzed using the HOLE2 software.^46^

### Cell culture for electrophysiological measurements

HEK293T cells (CLS Cat# 300192/p777_HEK293, RRID:CVCL_0045) were grown in DMEM culture medium and were maintained at 37 °C with 10% CO_2_.

For 96-well-based calcium assays, HEK293T cells grown in T25 flasks were co-transfected with 6 µg of wild-type or mutant hTRPM3-GFP DNA cloned in the pCAGGSM2 vector and 6 µg of the genetically encoded calcium indicator jRCAMP1b, using TransIT-LT1 transfection reagent (Mirus Bio, Madison, WI, USA). Following a 24-h incubation, cells were harvested and seeded at 100,000 cells/well into Poly-L-Lysine–coated 96-well black-wall/clear-bottom plates (Greiner Bio One, Frickenhausen, Germany), and cultured overnight before fluorescence recording.

For single-cell imaging and electrophysiological experiments, cells were grown in six-well plates and, upon reaching ∼50% confluency, transfected with 2 µg of wild-type or mutant hTRPM3-GFP construct using TransIT-LT1 transfection reagent followed by overnight incubation. Cells where then harvested and reseeded on 24-mm glass coverslips for use within 4-8 hours.

### Well-based calcium imaging

To obtain concentration–response curves for ligand action at wild type and mutant TRPM3, 96-well plated containing HEK293T-cells co-expressing the channel and jRCAMP1b were washed and filled with extracellular solution containing 150 NaCl, 4 KCl, 2 CaCl_2_, 2 MgCl_2_ and 10 HEPES (pH 7.4 with NaOH), and placed in the Hamamatsu FDSS/µCell kinetic plate imager. jRCAMP1b fluorescence (excitation: 560 nm; emission: 590 nm) was monitored at 0.5 Hz, while TRPM3 ligands at a range of concentrations were automatically pipetted into the wells from preloaded compound plates. At the end of the experiments, all wells were stimulated with a solution containing the calcium ionophore ionomycin (to a final concentration of 20 μM) and 20 mM CaCl_2_, to produce saturation of the jRCAMP1b fluorescence for normalization. All data are expressed as ΔF/F_0,Normalized_, where ΔF represents the change from baseline and F_0,Normalized_ the normalized baseline fluorescence. Experiments were performed at room temperature (22 ± 2°C).

### Cell-based calcium imaging

Changes in intracellular calcium concentration were monitored using ratiometric Fura-2-based fluorimetry. Cells were loaded with 2 μM Fura-2-acetoxymethyl ester (ION Biosciences) for 30 min at 37 °C. Fluorescence was measured during alternating illumination at 340 and 380 nm using either a Cell^M^ (Olympus) or Eclipse Ti (Nikon) fluorescence microscopy system, and absolute calcium concentrations were calculated from the ratio of the fluorescence signals at these two wavelengths (R = F_34_0/F_380_) as [Ca^2+^]_i_ = K_m_ × (R − R_min_)/(R_max_ − R), where K_m_, R_min_ and R_max_ were estimated based on calibration experiments with known calcium concentrations. The standard extracellular solution contained (in mM): 150 NaCl, 4 KCl, 2 CaCl_2_, 2 MgCl_2_ and 10 HEPES (pH 7.4 with NaOH). When indicated, the standard extracellular solution was rapidly exchanged by solution containing the indicated concentration of TRPM3 (ant)agonists via a gravity-driven perfusion system.

### Data analysis and statistics

Data analysis was performed using custom-written routines in Igor Pro (v.9; Wavemetrics); figure assembly and statistical testing were performed using Origin 2023B (OriginLab) and. One-way ANOVA with a Tukey post-hoc test was used to compare the effect of multiple mutations on TRPM3 ligand responses. P < 0.05 was considered as statistically significant.

### Materials

Pregnenolone sulfate (PS), primidone, ononetin, and ionomycin were obtained from Sigma-Aldrich. Isosakuranetin was purchased from Extrasynthese, Merck, Carl Roth GmbH and Sigma. The Chemical Abstracts Service Registry number provided by Extrasynthese, Merck and Carl Roth GmbH was CAS No. 480-43-3, which refers to (*2S*)-isosakuranetin. Sigma-Aldrich provides plant-derived isosakuranetin with CAS No. 26207-61-4, which indicates no specific stereochemistry in the structure. *(2S)-* isosakuranetin and *(2R)*-isosakuranetin were separated from racemic isosakuranetin using chiral HPLC. Racemic CIM0216 was obtained from Tocris, and separated using chiral HPLC to obtain pure *(R)*-CIM0216 and *(S)*-CIM0216.

### Chiral separation of racemic mixtures

The chirality of commercially acquired isosakuranetin was analysed with a Waters Acquity UPLC H-Class UPLC system and was a ChiralPack IG (150 x 4.6 mm)x3 µm, chiral column. Methanol was used as isocratic eluent with a flow of 1ml/min. UV detection was performed with an ACQUITY UPLC PDA eLambda Detector (200-420 nm).

Purification of (S)- and (R)-isosakuranetin for functional studies was carried out on an HPLC system combining a Waters 2489 UV/Visible Detector, a Waters 2545 Binary Gradient Module, a System Fluidics Organizer, a 515 HPLC Pump, a Waters 2767 Sample Manager and a Waters MS3100 Mass detector. The chiral column used was a ChiralPack IG (250 x 20 mm)x5 µm using methanol as isocratic eluent with a flow of 20ml/min. UV detection was performed at 290 nm. The optical rotation of both peaks was measured using a polarimeter (KRUSS P3000). The second peak has an [α]_D_ of -70°, in agreement with the reported data ^47^.

Chiral resolution of CIM0216 was performed on a Waters Thar SFC-80 system composed of a binary pump for delivering CO_2_ and modifier, an injection module, a diode array detector and a fraction collection module. The chiral column used was a (R,R)Whelk -01 (250 x 4.6 mm)x5 µm using 25% methanol and 75% CO_2_ as isocratic eluent with a flow of 3g/min (Back pressure : 100 bar and Temperature : 30°C). UV detection was performed at 214.0 nm. The absolute configuration of the two separated enantiomers was determined via Vibrational Circular Dichroism (VCD) analysis. Each enantiomer was dissolved in CDCl3 (10 mg in 50 µL), and the solution was transferred to a BaF_2_ cell. VCD spectra were acquired using a JASCO FVS-6000 instrument. The absolute configurations were assigned by comparing the experimentally measured VCD spectrum of each enantiomer with computationally predicted spectra.

## Data availability

Atomic models have been deposited in the Protein Data Bank (PDB), and cryo-EM maps, corresponding half maps and masks can be found at the Electron Microscopy Data Bank (EMDB). The atomic models have been deposited with accession codes 9QHN (TRPM3_APO_), 9QHO (TRPM3_PRM_), 9QHP (TRPM3_ISO_), 9QHM (TRPM3_ONO_) and 9QHQ (TRPM3_AGO_ ; in the combined presence of R-CIM0216, PS and PI(4,5)P_2_). The corresponding maps can be accessed under EMD-53174 (TRPM3_APO_), EMD-53175 (TRPM3_PRM_), EMD-53176 (TRPM3_ISO_), EMD-53173 (TRPM3_ONO_) and EMD-53177 (TRPM3_AGO_).

---

**Supplementary Figure 1.**
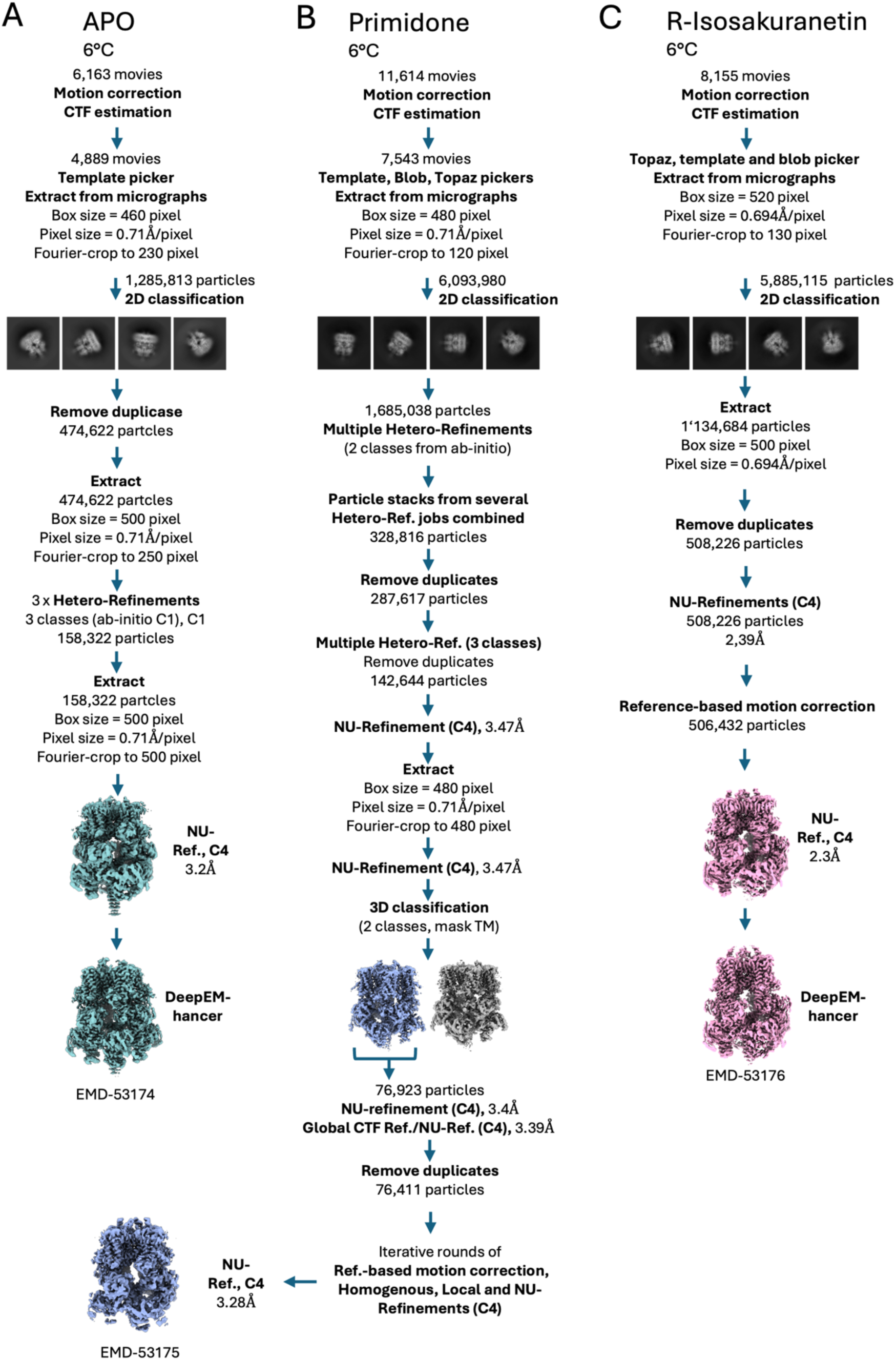
Cryo-EM workflow for TRPM3 in the apo-state (A), with primidone (B) and R-isosakuranetin (C).

**Supplementary Figure 2.**
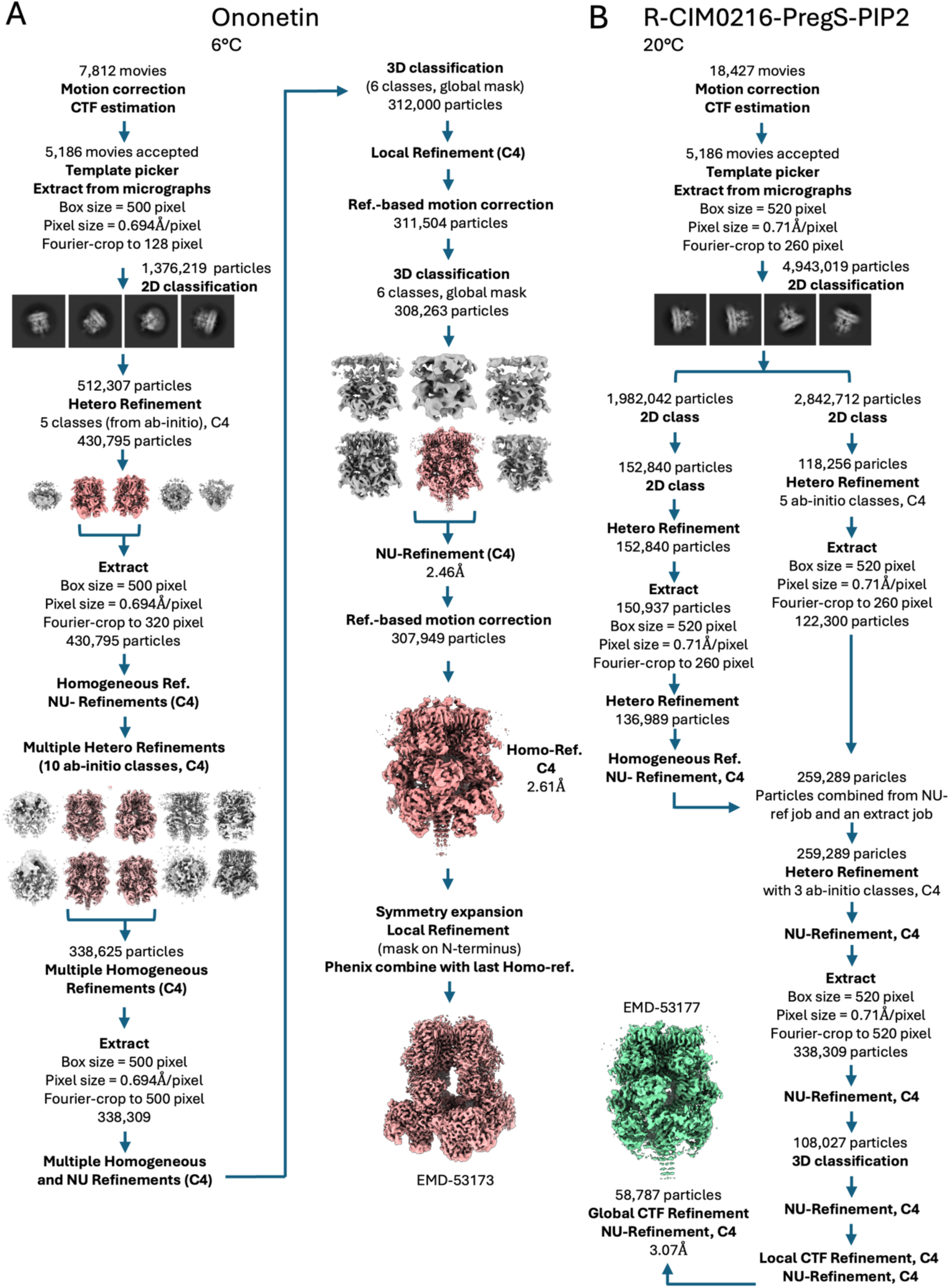
Cryo-EM workflow for TRPM3 with ononetin (A) and R-CIM0216 (B).

**Supplementary Figure 3.**
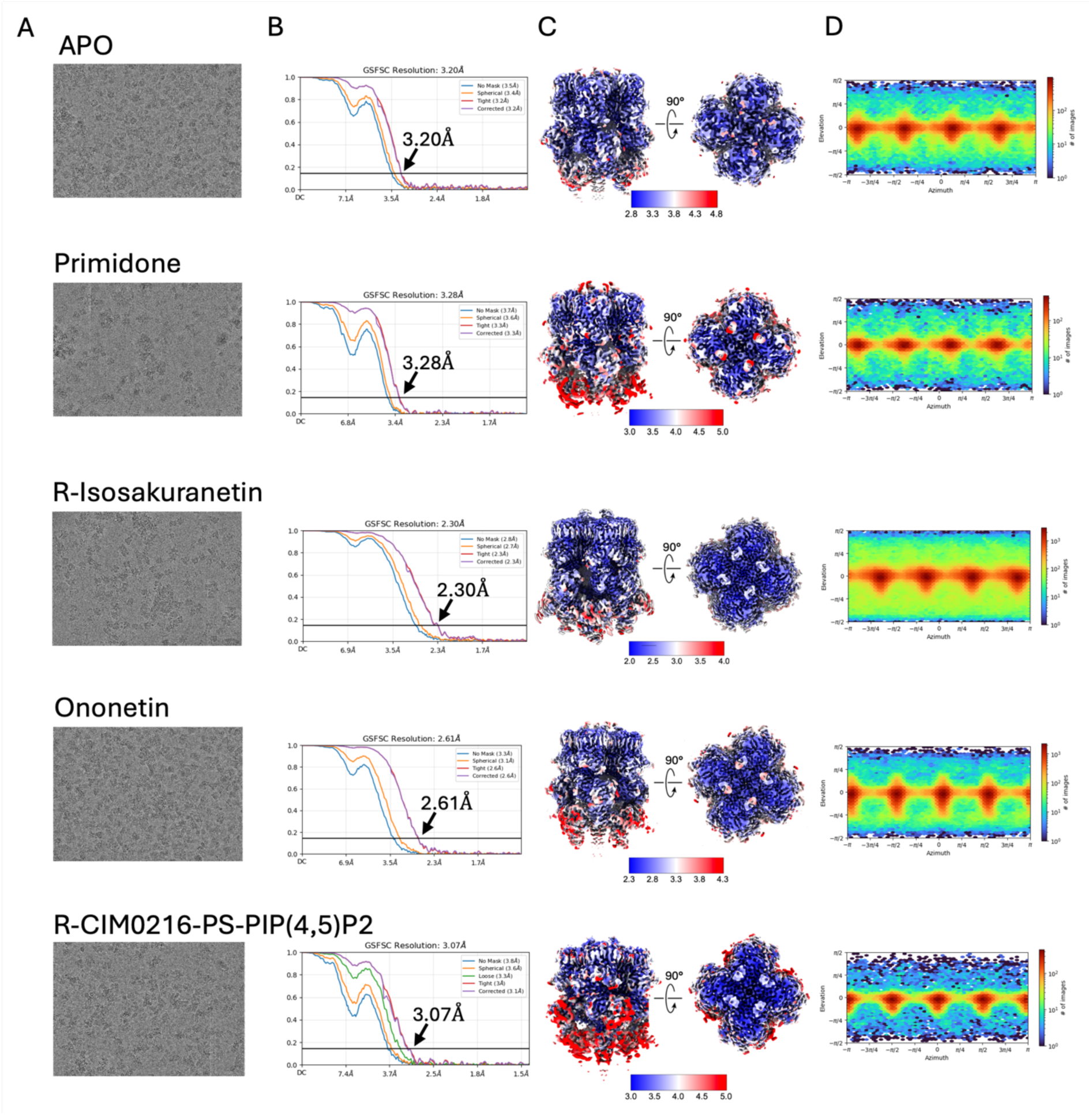
Local resolution and Fourier shell correlaton plots of TRPM3 in the apo state or with ligands. (A) Representative micrographs. (B) Fourier shell correlation (FSC) curves of the final reconstructions with the resolution indicated in Å (marked at the 0.143 criterion). (C) Filtered maps shown from the side and from extracellular (color-coded for local resoluton). (D) Angular distribution plot of particles.

**Supplementary Figure 4.**
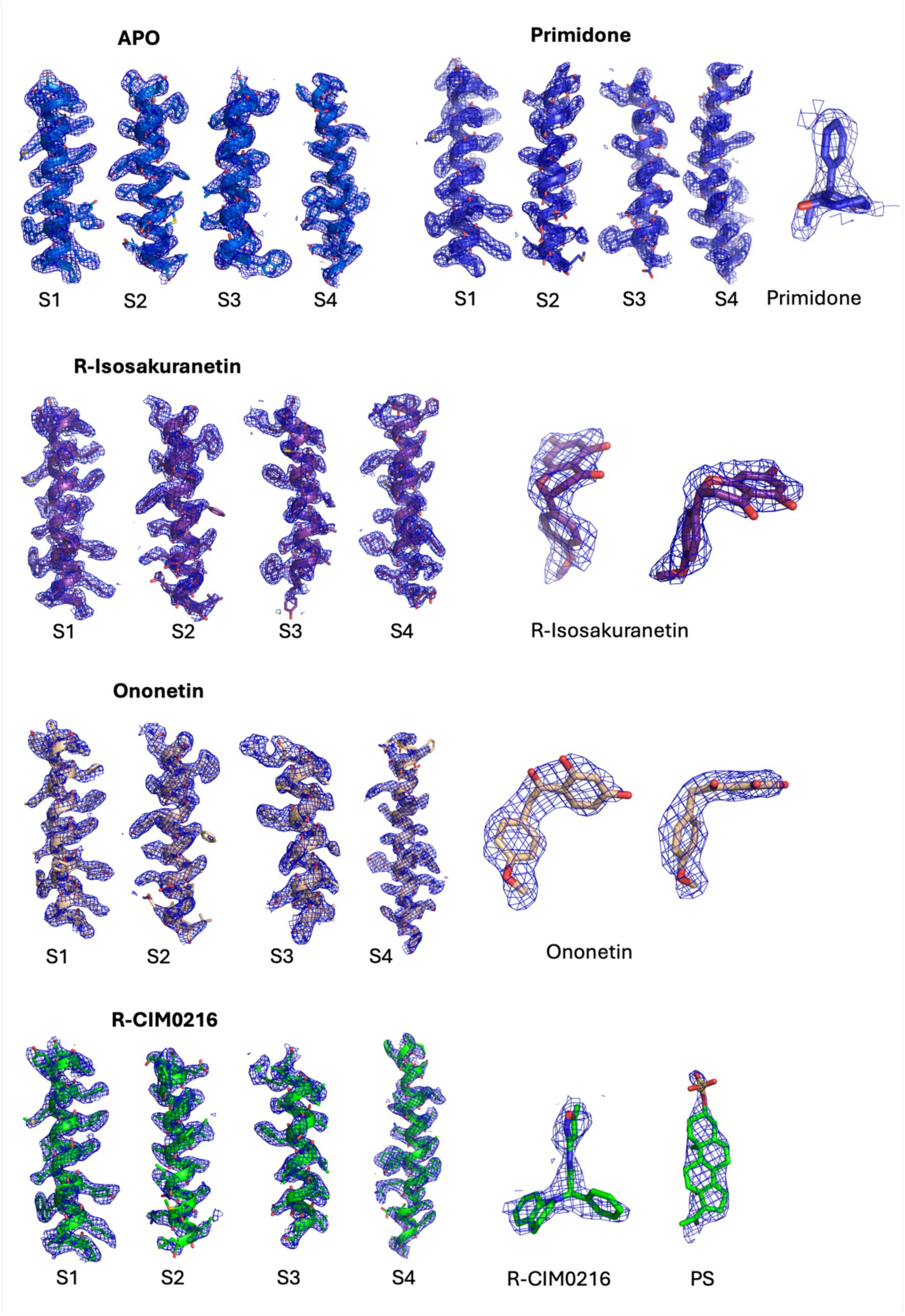
Map and model quality. Agreement of the cryo-EM density map with the final model shown for the S1-S4 TM-helices of TRPM3 in the apo-state or with ligands. The map is contoured at between 4-7σ for the apo-state, 8-10 σ for primidone, 7-10 σ for R-isosakuranetin, 4-8 σ for ononetin and 7-9 σ for R-CIM0216 in pymol.

**Supplementary Figure 5.**
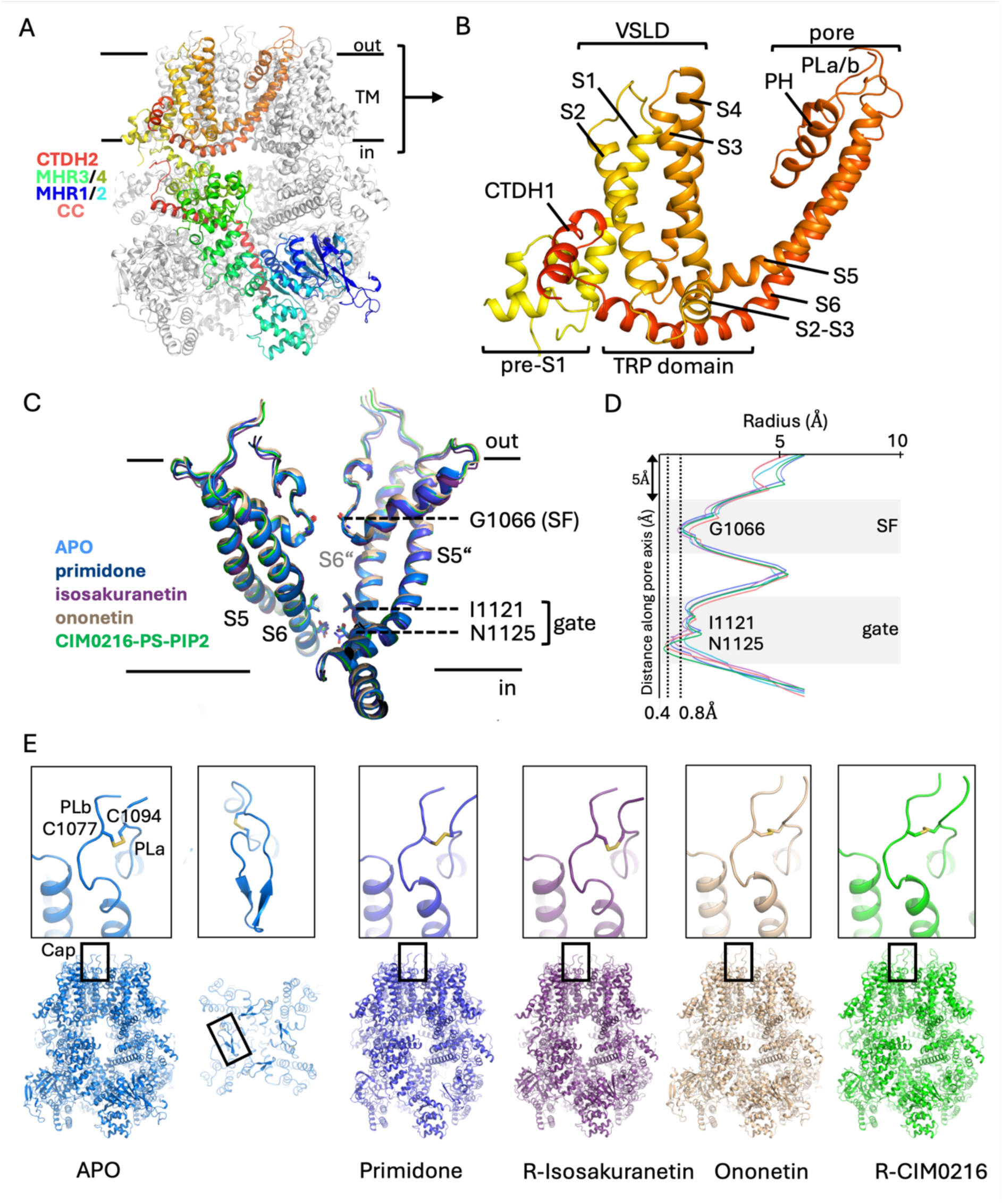
TRPM3 topology. (A) Architecture of TRPM3 with one chain displayed in rainbow colours (N-terminal blue, C-terminal red). (B) Enlarged view of (A) showing the organization of the TM domain in cartoon representation and with helices labelled. VSLD; voltage sensor like domain, CTDH; C-terminal domain helix, PH; pore helix (C) View from the side on the pore with the pore forming helices and important residues labelled. SF; selectivity filter, respectively. (D) HOLE analysis indicating close apposition of G1066, I1121 and N1125 and non-conductive conformations for all structures obtained in this study. Color code as in (C). (E) TRPM3 cap region and close-up view on a disulfide bridge with critical role during activation. The full cap region is shown according to an AlphaFold3 ^48^ prediction of TRPM3. In (C), (D) and (E) the human numbering of residues is shown (corresponding to +10 residues compared to the numbering in mouse TRPM3a2).

**Supplementary Figure 6.**
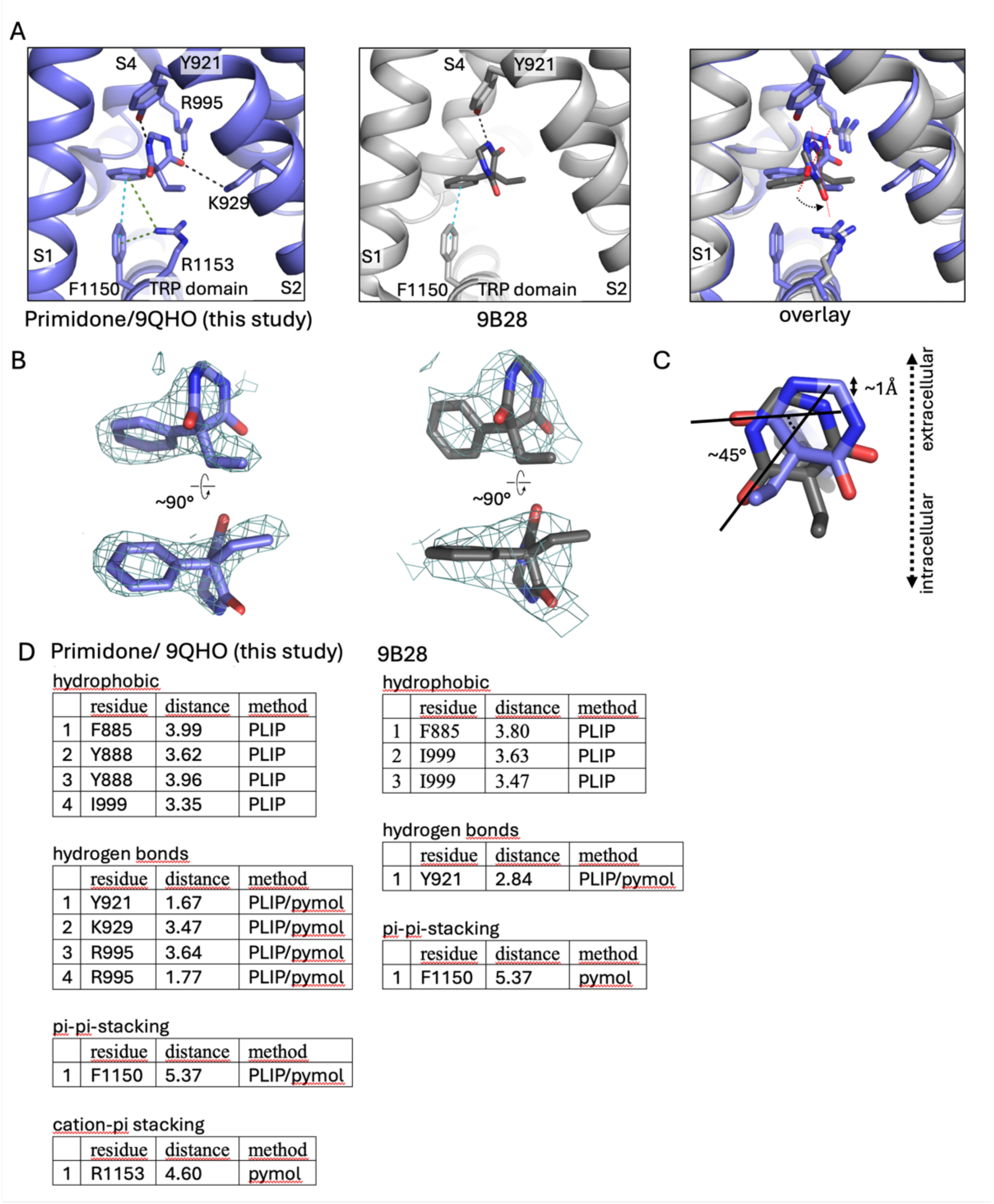
The primidone binding site. (A) Close-up view on the primidone binding site with residues and ligands shown in cartoon representation and as sticks. Direct interactions are highlighted as dashed lines; hydrogen bonds in black, cation-π and π-π stacks in pink and cyan, respectively. Left: this study/9QHO, middle: Yin et al./9B28, right: overlay. (B) Primidone and corresponding cryo-EM densities are shown. Maps contoured at 9 σ (this study, 9QHO) and 7 σ for 9B28, respectively. In (A) and (B) only direct interactions are shown. (C) Overlay of the binding poses of primidone in our study with primidone of 9B28, angular shift and dislocation indicated. (D) Analysis of the interactions with primidone (this study/9QHO vs. 9B28) using pymol and the Protein Ligand Interaction Profiler (PLIP). In (A) and (C) the human numbering of residues is shown.

**Supplementary Figure 7.**
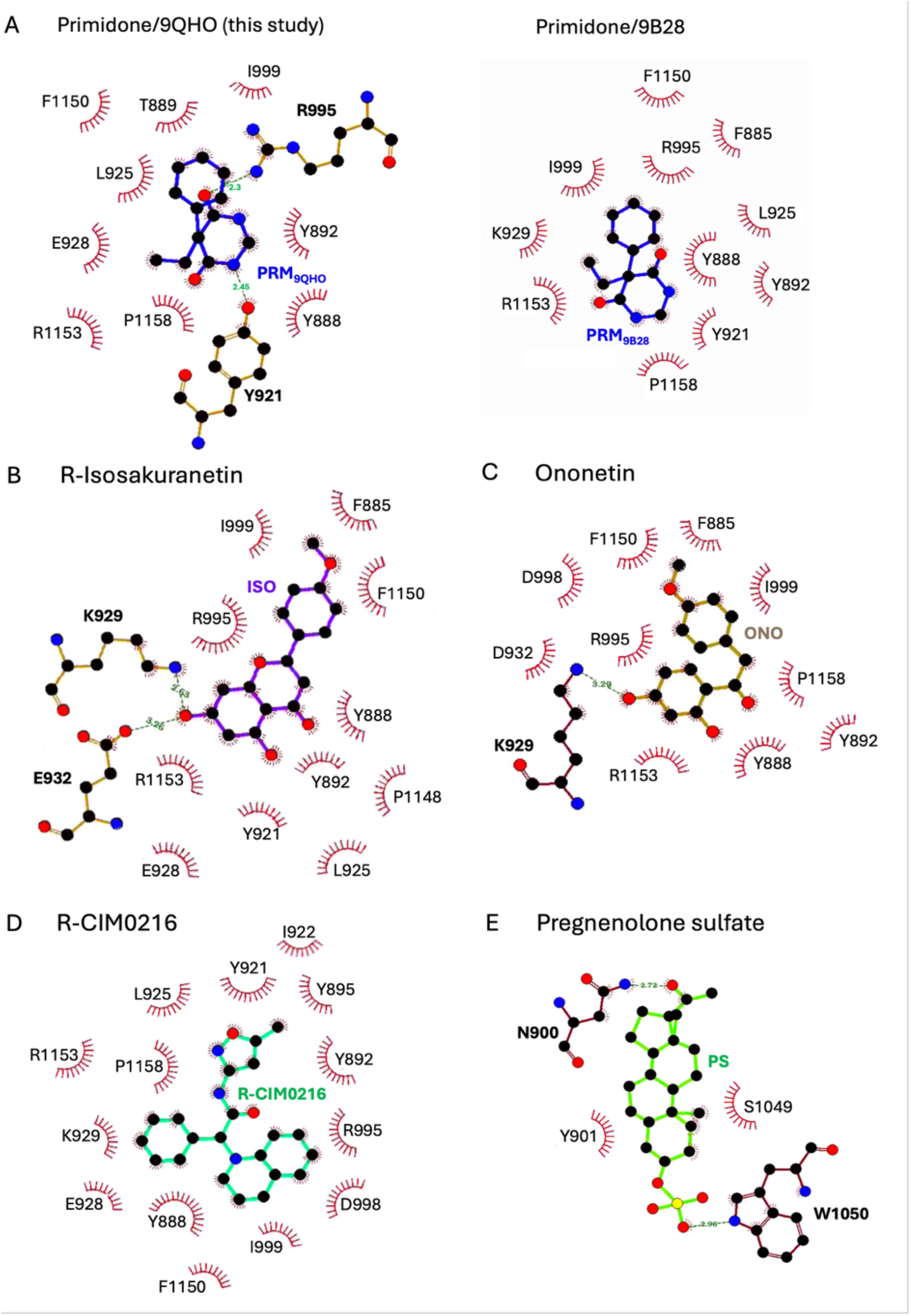
LigPlot+ analysis of the residues interacting with ligands. (A) primidone, (B) R-isosakuranetin, (C) ononetin, (D) R-CIM0216 and (E) PS. The ligands and protein side chains are shown in ball-and-stick representation, with the ligand bonds coloured in blue for primidone, violet for R-isosakuranetin, wheat for ononetin and green for R-CIM0216. Hydrogen bonds are indicated as green dashed lines, while the spoked arcs represent protein residues that form non-bonded contacts with the ligand (π-cation stacks and hydrophobic contacts including π-π stack interactions).

**Supplementary Figure 8.**
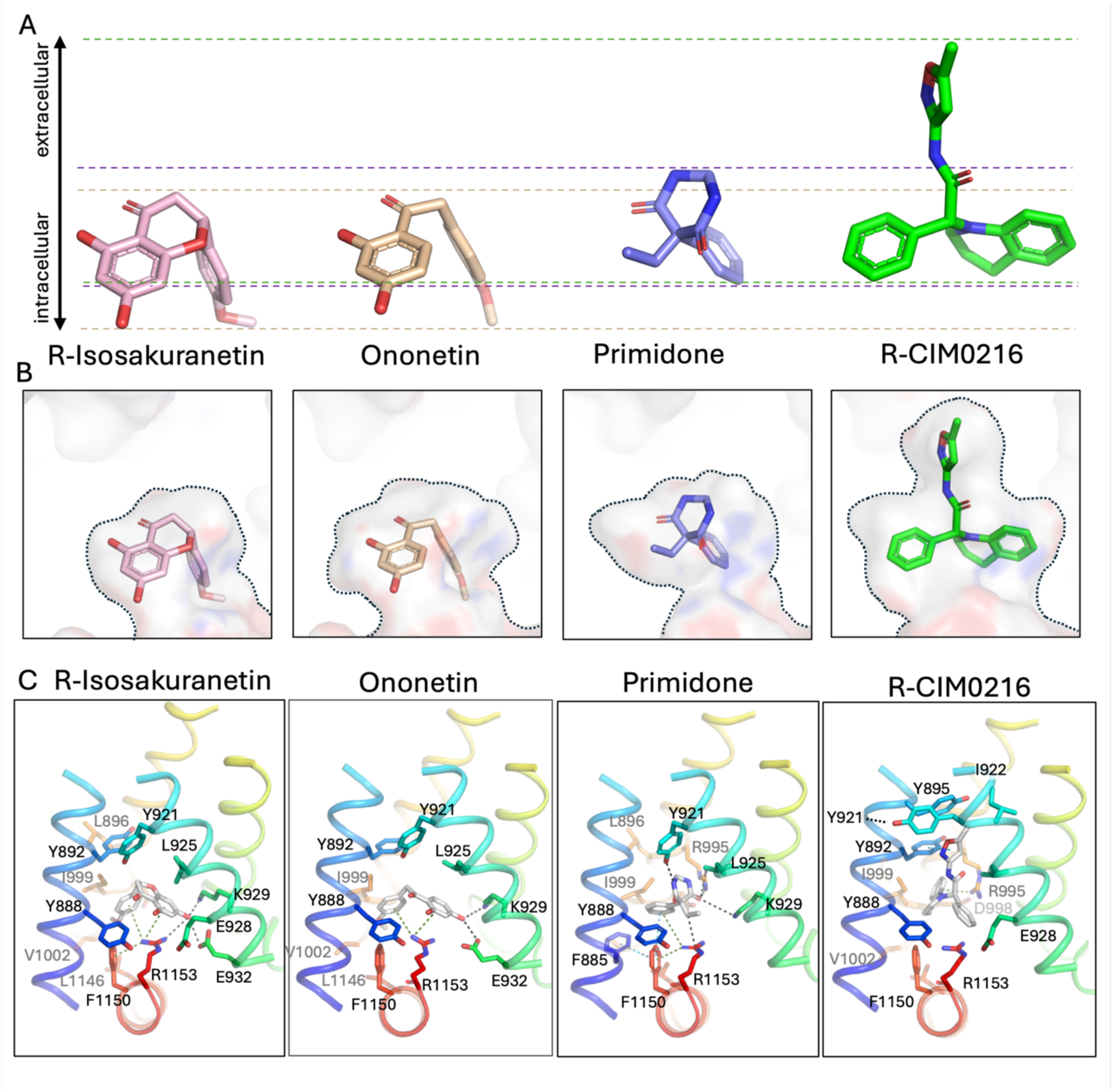
Relative position of antagonists and agonists and interactions inside the ligand binding pocket. (A) Relative position of the ligands along the intracellular-extracellular axis. Upper and lower boundaries of the ligands are marked with dashed lines in the coloration of the respective ligand. R-CIM0216 extends substantially farther to the extracellular side compared to the antagonists. (B) Outline of the ligand binding pockets for a comparison of the occupied space by the respective ligands. (C) Close-up view on the ligand binding site. S1-S4 helices and TRP domain are represented as cartoon model with residues shown as sticks that directly interact with the ligands R-isosakuranetin, ononetin, primidone or R-CIM0216. The atomic models are shown in rainbow colors to highlight similarities and distinctions regarding the ligand interactions with respective residues. Ligands are coloured in grey and the human numbering of residues is shown.

**Supplementary Figure 9.**
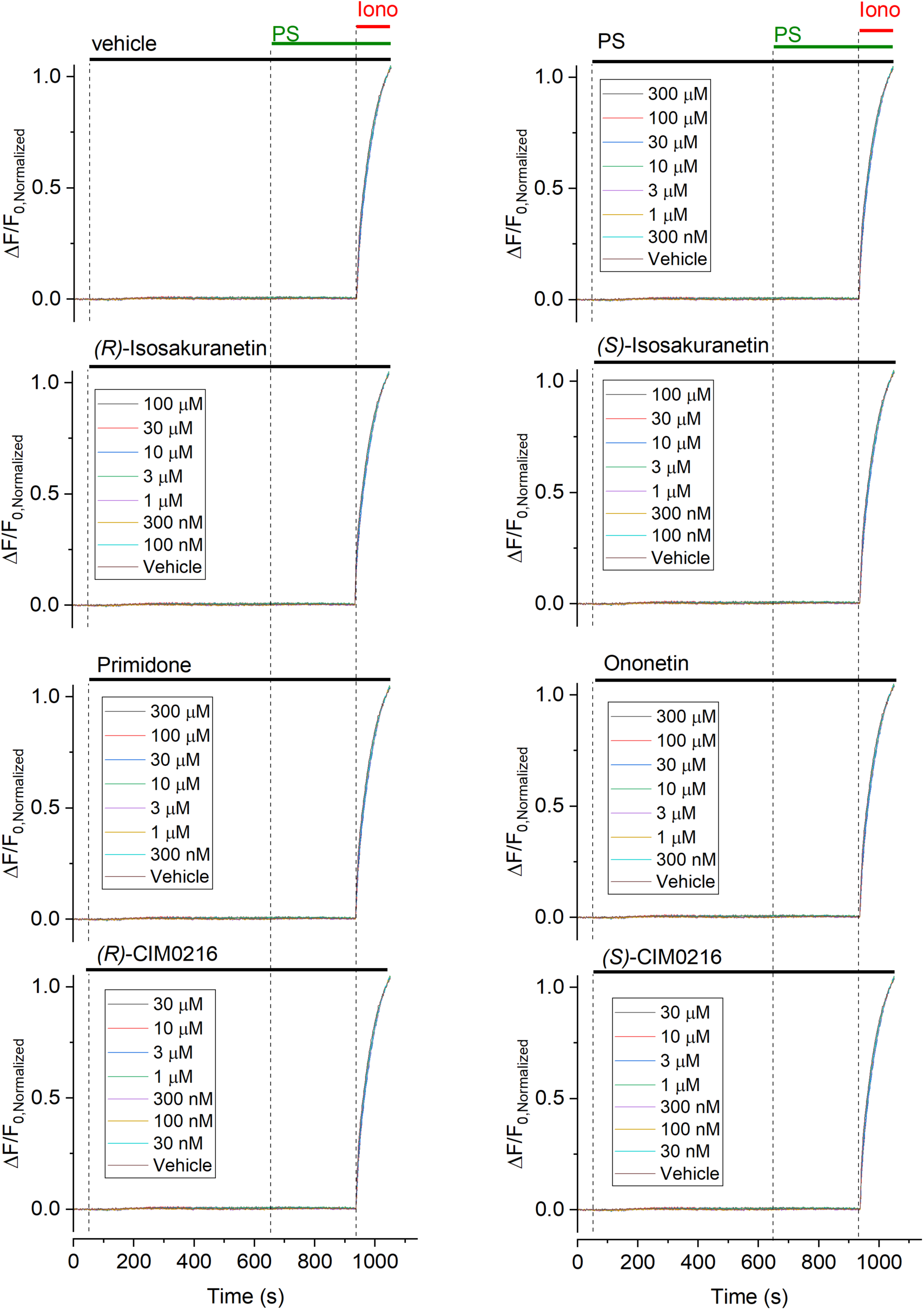
Lack of response to TRPM3 ligands in HEK293 cells expressing jRCaMP1b but not TRPM3. HEK293 cells transfected with jRCaMP1b were pre-incubated with vehicle or with the indicated concentrations PS, (R)- and (S)-isosakuranetin, primidone, ononetin, (R)-CIM0216 and (S)-CIM0216, followed by a test stimulus using PS (50 μM). Finally, cells were exposed to the ionophore ionomycin in the presence of excess extracellular Ca^2+^ (20 mM) to obtain saturation of the jRCaMP1b signal. No detectable changes in jRCaMP1b were observed for any of the TRPM3 ligands.

**Supplementary 10.**
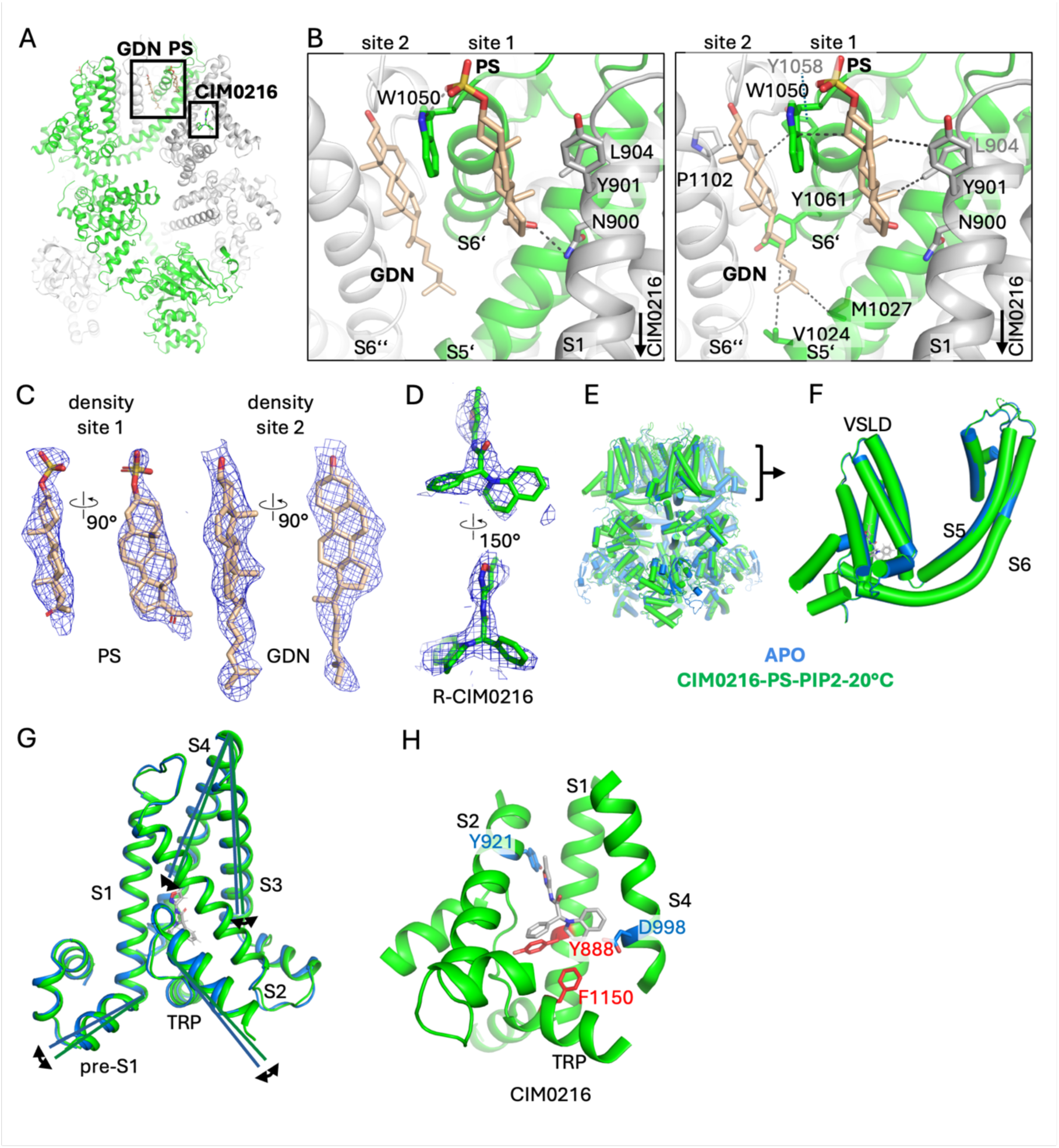
PS and CIM0216 binding sites. (A) TRPM3 tetrameric model with agonist binding sites emphasized by boxes. Visible chains are coloured in grey, green and white. (B) Binding sites of PS (site 1) and a glycodiosgenin (GDN) detergent molecule at the potential PS binding site 2 sandwiched between helices S1, S5’, S6’ and S6’’. Interactions are shown as dashed lines and corresponding residues as sticks for hydrophilic interactions (left) and hydrophobic interactions (right). Human numbering of residues is shown. PS and GDN (C) and R-CIM0216 (D) with corresponding cryo-EM densities contoured between 6 and 7 σ in pymol. (E) Structural alignment of TRPM3 tetramers in the apo-state (marine) and bound to agonists (green) with helices shown as cylinders. (F) close-up view of F on TM part. (G) Close-up view on S1-S4 with helices represented in cartoon format. Lines and arrows indicate conformational changes accompanied by agonist binding. Colours as in (E,F) (H) Close-up view on S1-S4 with helices represented in cartoon format. Residues that turn the CIM0216 agonist into an antagonist (red) or make it more potent (blue) are shown.

**Supplementary Figure 11.**
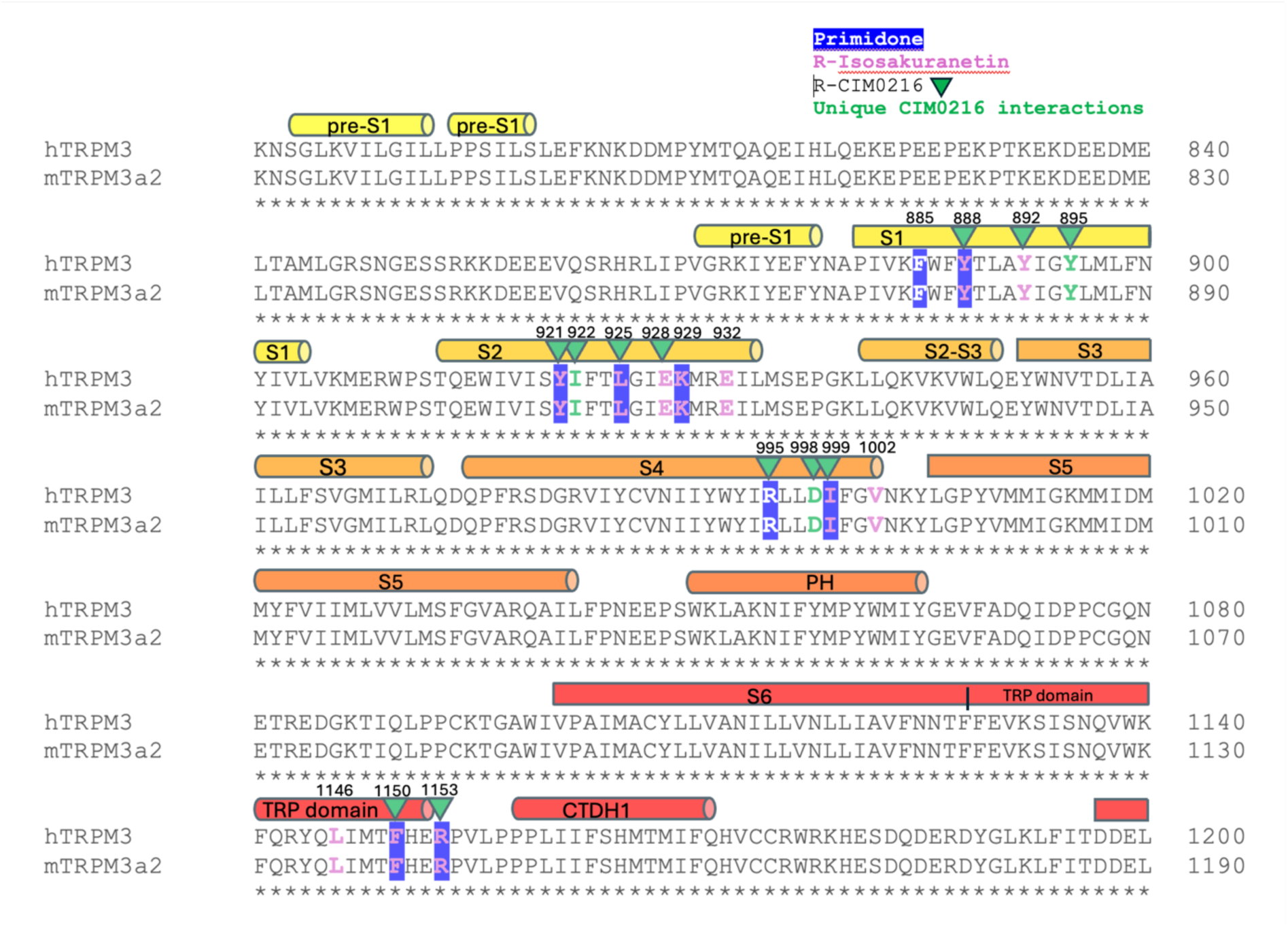
Sequence alignment. Sequence alignment of the TM region between human and mouse TRPM3a2 isoforms. Helices are shown as cylinders, labelled and coloured like in (A) and (B). Residues that interact with primidone, *(R)*-isosakuranetin or *(R)*-CIM0216 are marked as blue boxes around letters, as violet letters or indicated by green triangles, respectively, with the numbering of the human protein given

**Supplementary Figure 12:**
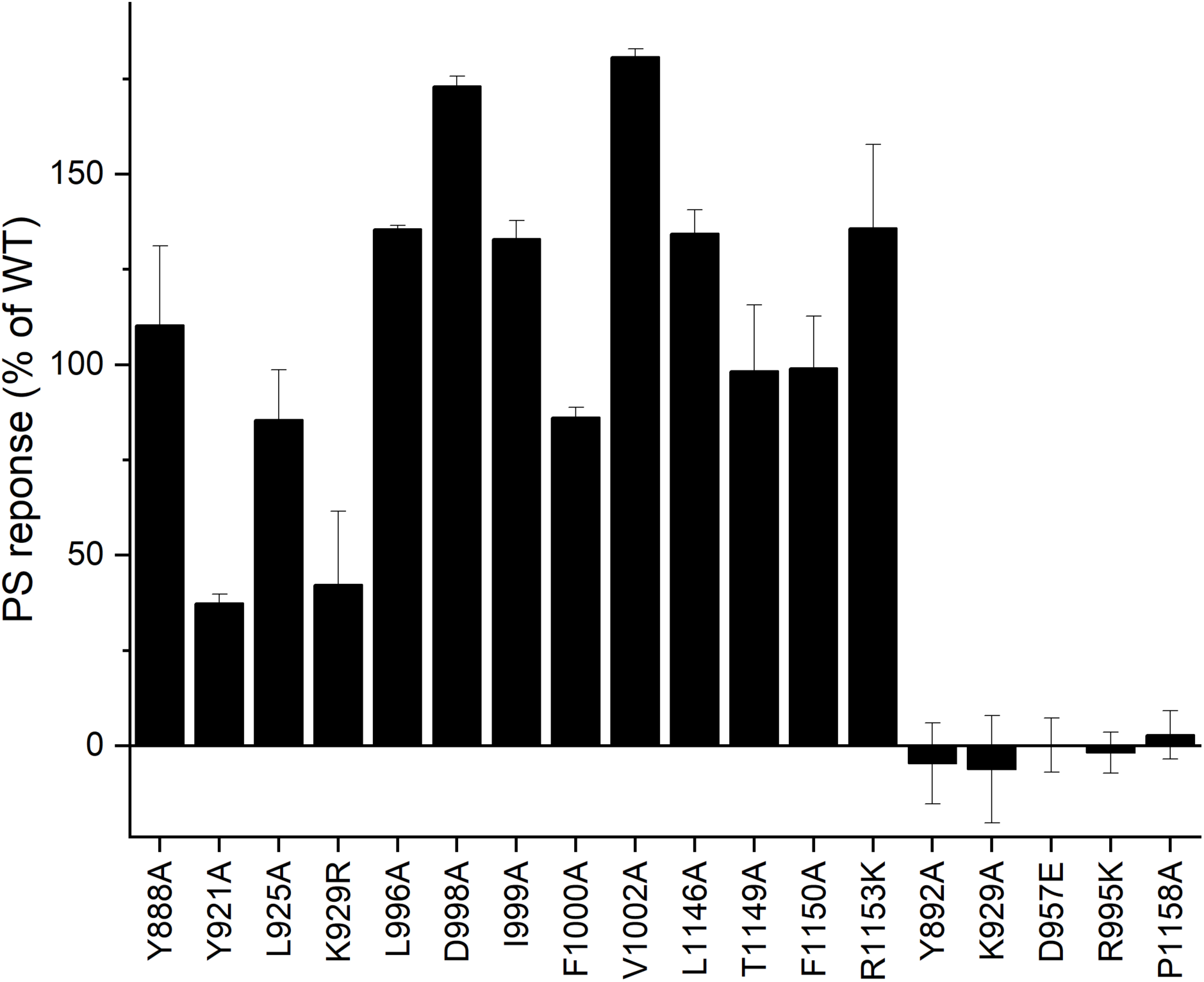
Functional and non-functional mutants. Summary of calcium responses to PS (50 μM) measured in Fura-2-loaded cells expressing the indicated mutants. Responses were normalized to the mean PS response in cells expressing WT TRPM3 measured on the same recording day. Only mutants for which a PS response exceeding 25% of the WT response was measured were used in further experiments.

**Supplementary Figure 13:**
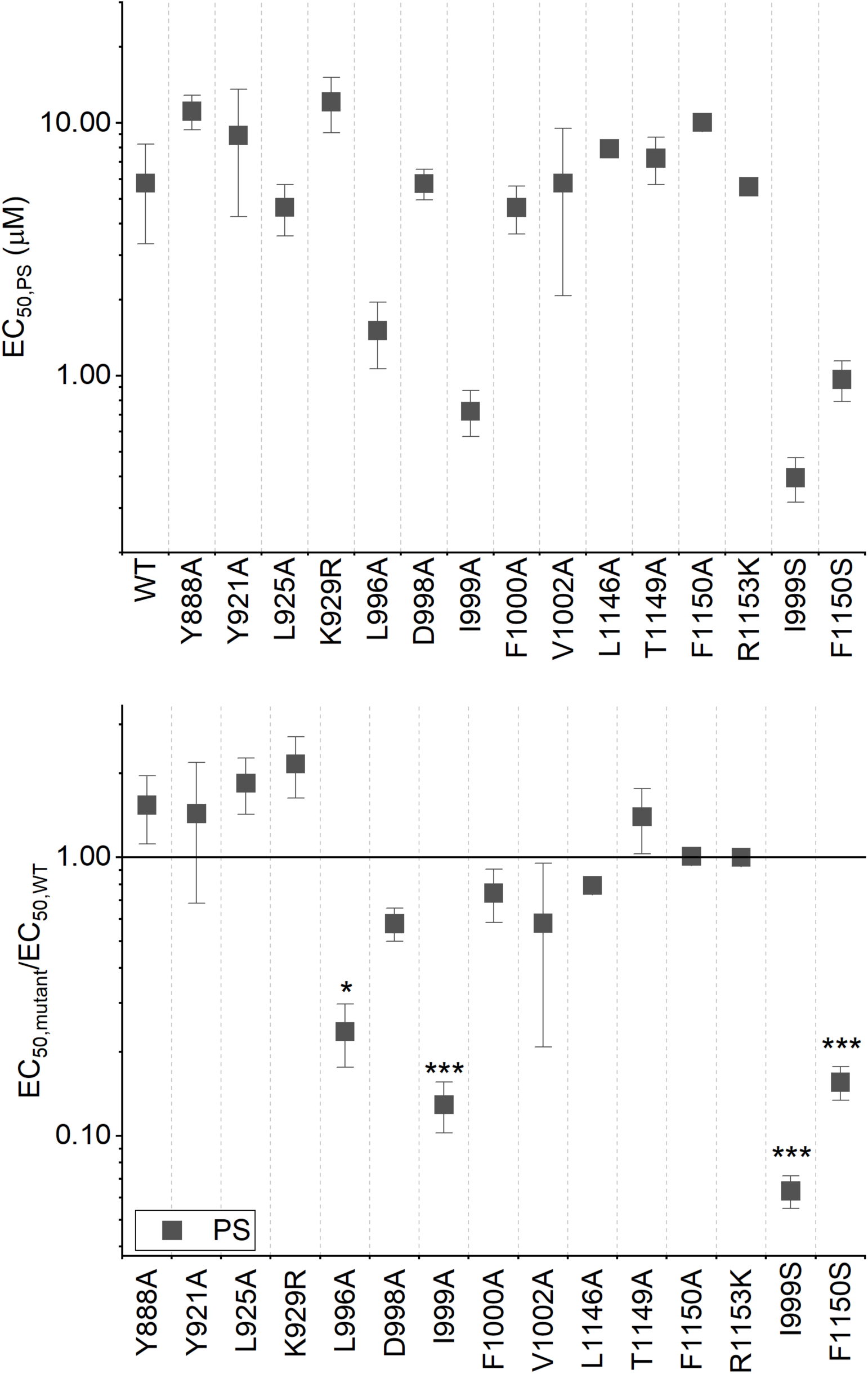
PS responses in WT and mutant TRPM3. (Top) EC_50_ values for the activation of WT and mutant TRPM3 by PS. (Bottom) EC_50_ values normalized to the values obtained for WT. *,***: p<0.05, 0.001 for the comparison with WT.

**Supplementary Figure 14.**
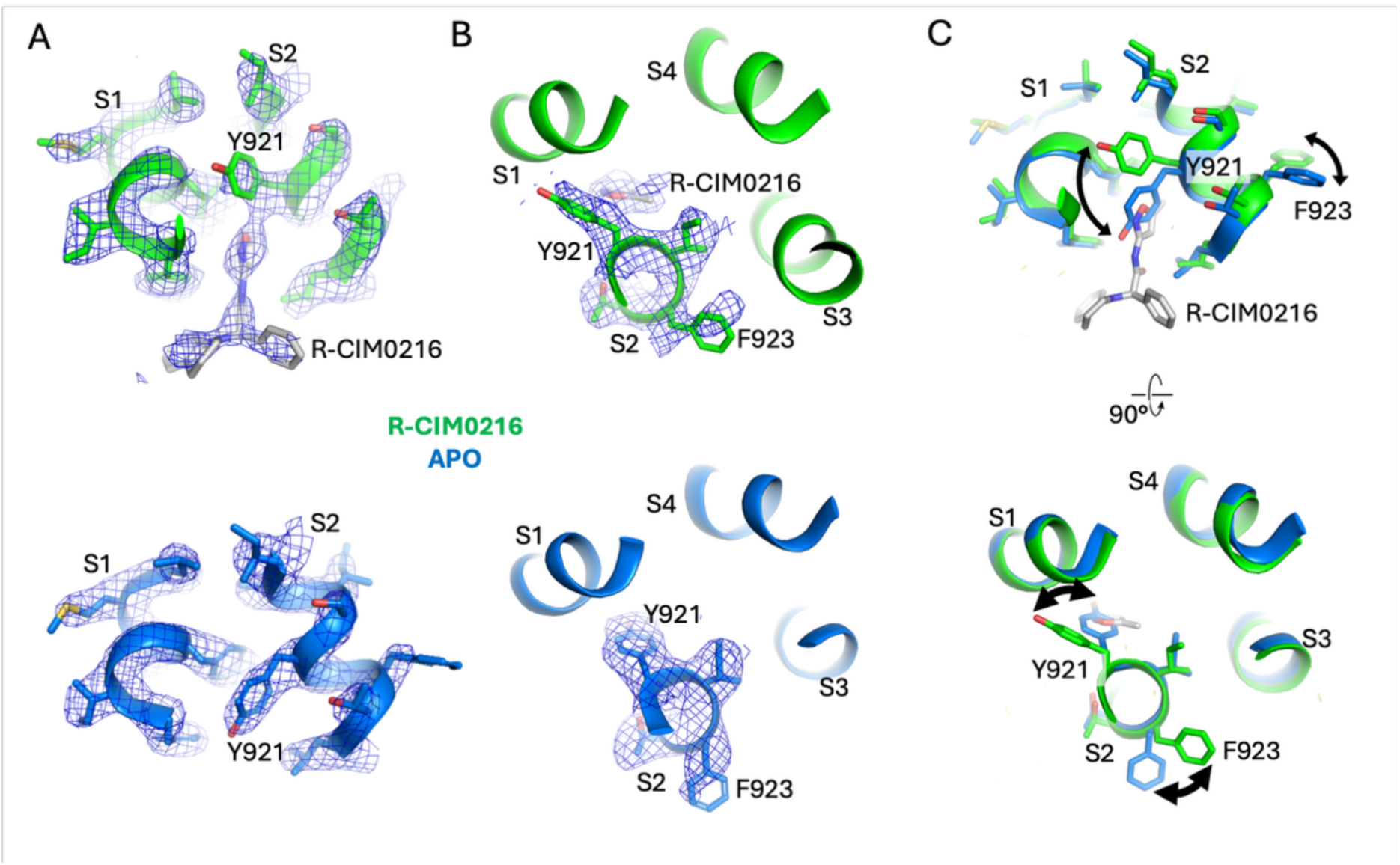
Conformational changes inside the ligand binding pocket associated with agonist binding. View from the side (A) or from extracellular (B) on the ligand binding site for R-CIM0216 compared to TRPM3 in the apo-state. The models are shown in cartoon representation with residues illustrated as sticks. Cryo-EM densities are overlaid and contoured between 9-11 σ in pymol. In case of R-CIM0216 an ‘upward’ movement of Y921 towards the extracellular side can be observed. (C) Overlay of atomic models in (A) and (B) to illustrate the conformational changes after agonist binding. The pose of Y921 observed in TRPM3_APO_ is also found for TRPM3_ONO_, TRPM3_ISO_ and TRPM3_PRM_. Human numbering of residues is shown.

**Supplementary Figure 15.**
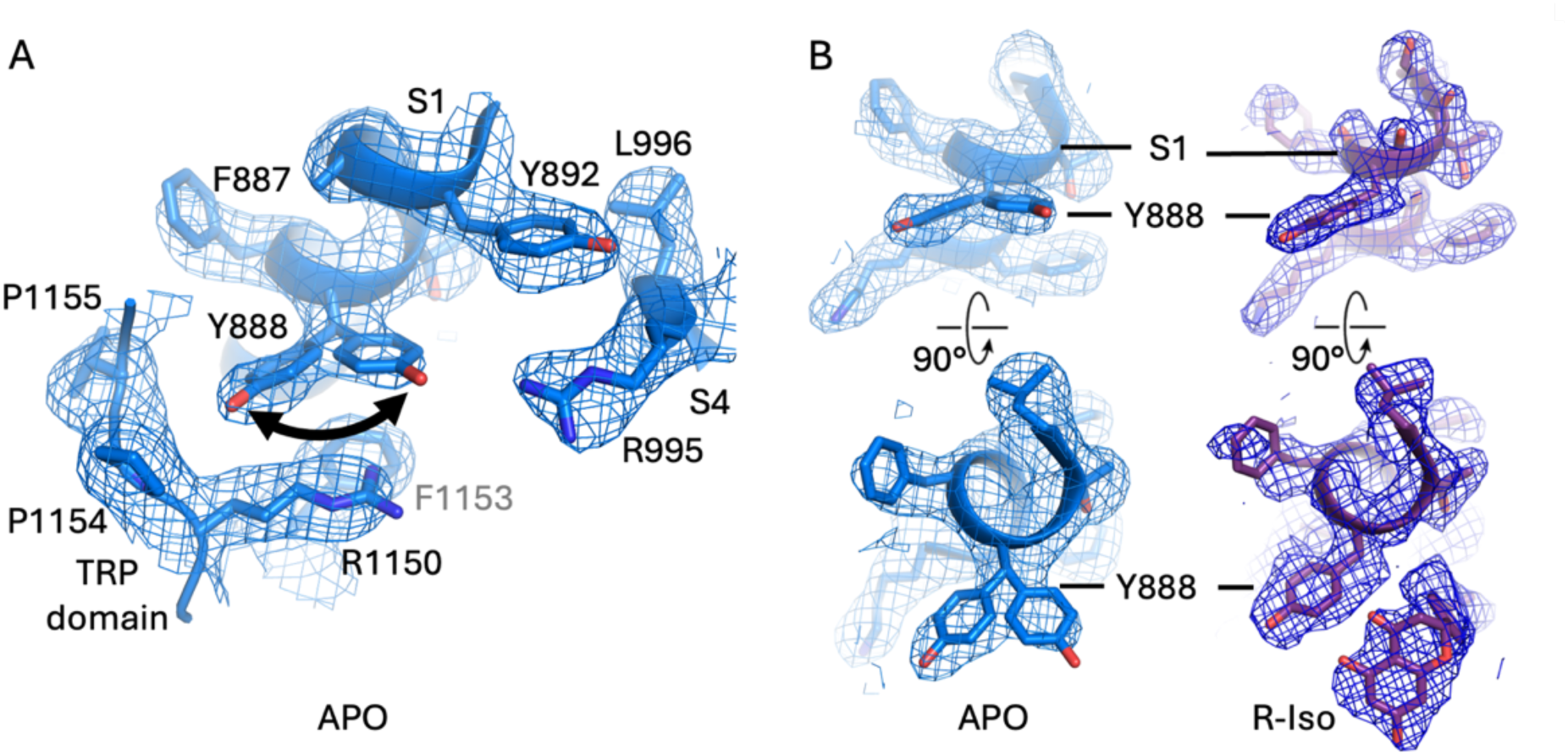
The position of Y888. (A) Illustration of the ligand binding pocket with view on Y888 in TRPM3 in the apo-state. In TRPM3_APO_ two rotamers can be observed as supported by the cryo-EM density. The atomic model is shown as cartoon with side chains displayed as sticks and labelled (B) Views from the side (top) and from extracellular (bottom) on Y888 in TRPM3_APO_ or bound to R-isosakuranetin, illustrating mobility of Y888 in the absence of bound ligands. The maps are contoured between 5.5 and 6 σ in pymol. The pose of Y888 observed for TRPM3_ISO_ is very similar in TRPM3_ONO_ and TRPM3_PRM_. The human numbering of residues is shown.

**Supplementary Figure 16.**
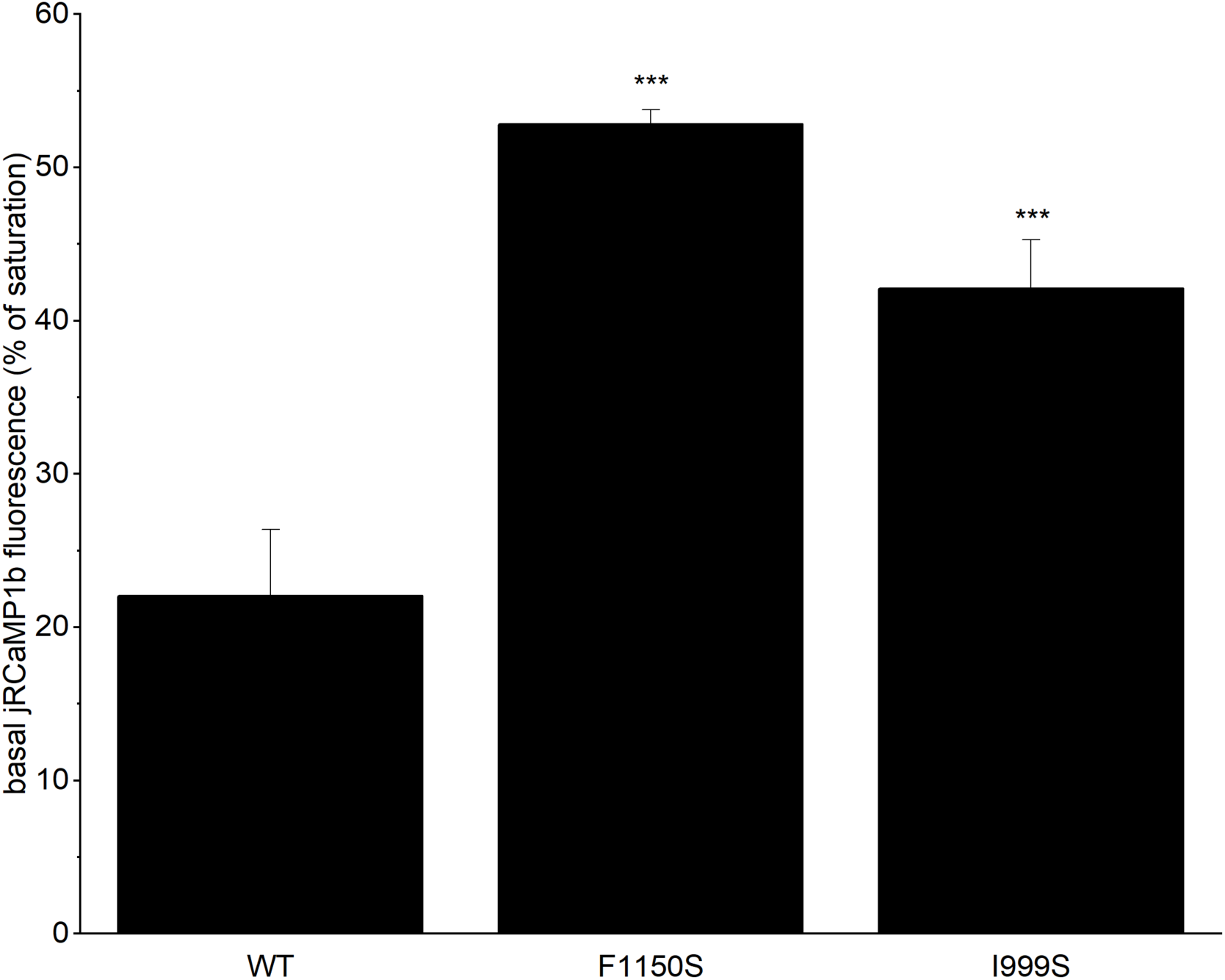
Increased basal calcium levels in HEK293 cells expressing the disease-associated mutants F1150S and IGGGS. Basal jRCaMP1b fluorescence was normalized to the maximal fluorescence upon final ionomycin stimulation, yielding a measure of non-stimulated cytosolic calcium levels in HEK293 cells expressing WT TRPM3 or the disease-associated mutation F1150S and I999S.

**Supplementary Table 1:**
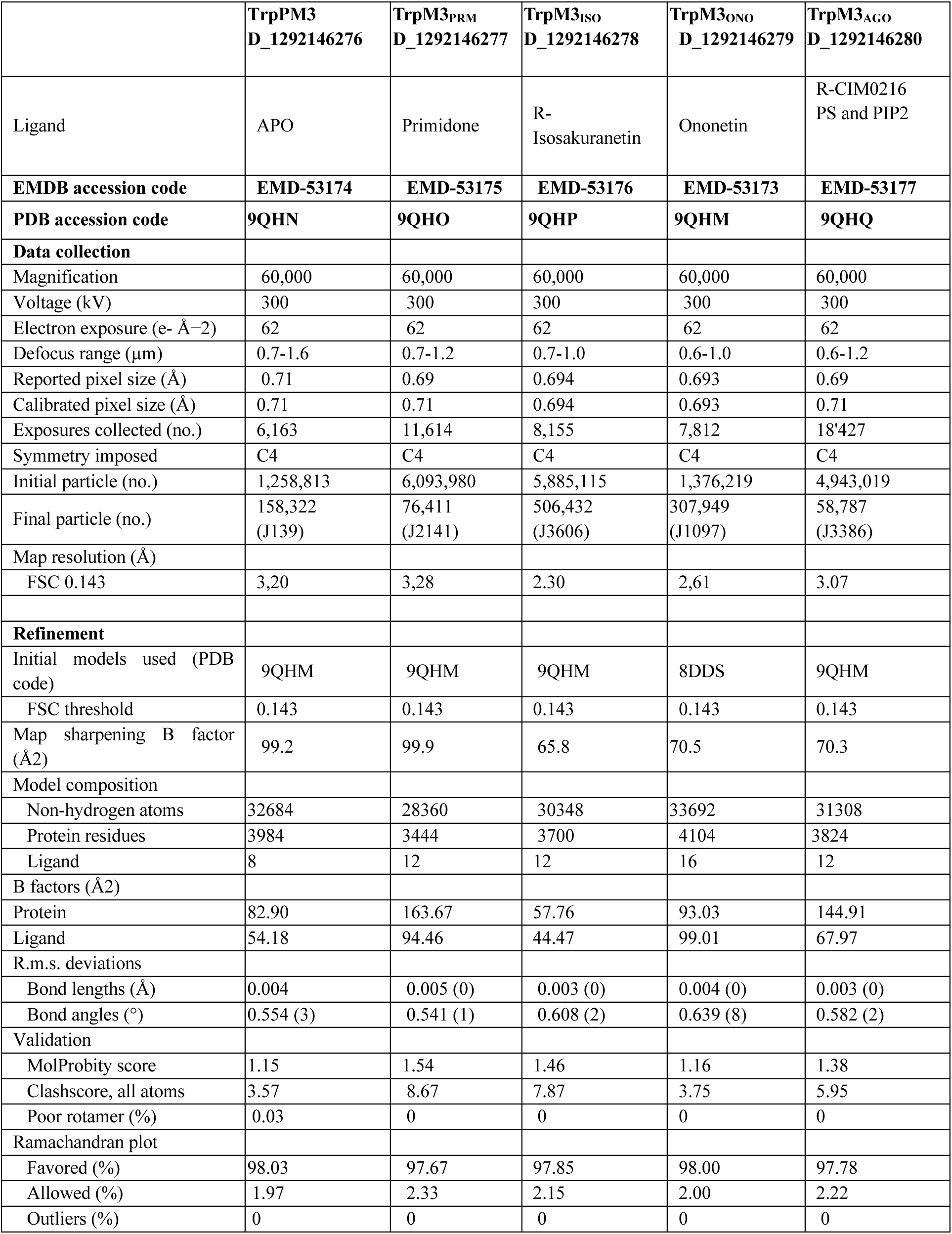
Cryo-EM data collection, refinement and validation statistics.

**Supplementary Table 2.**
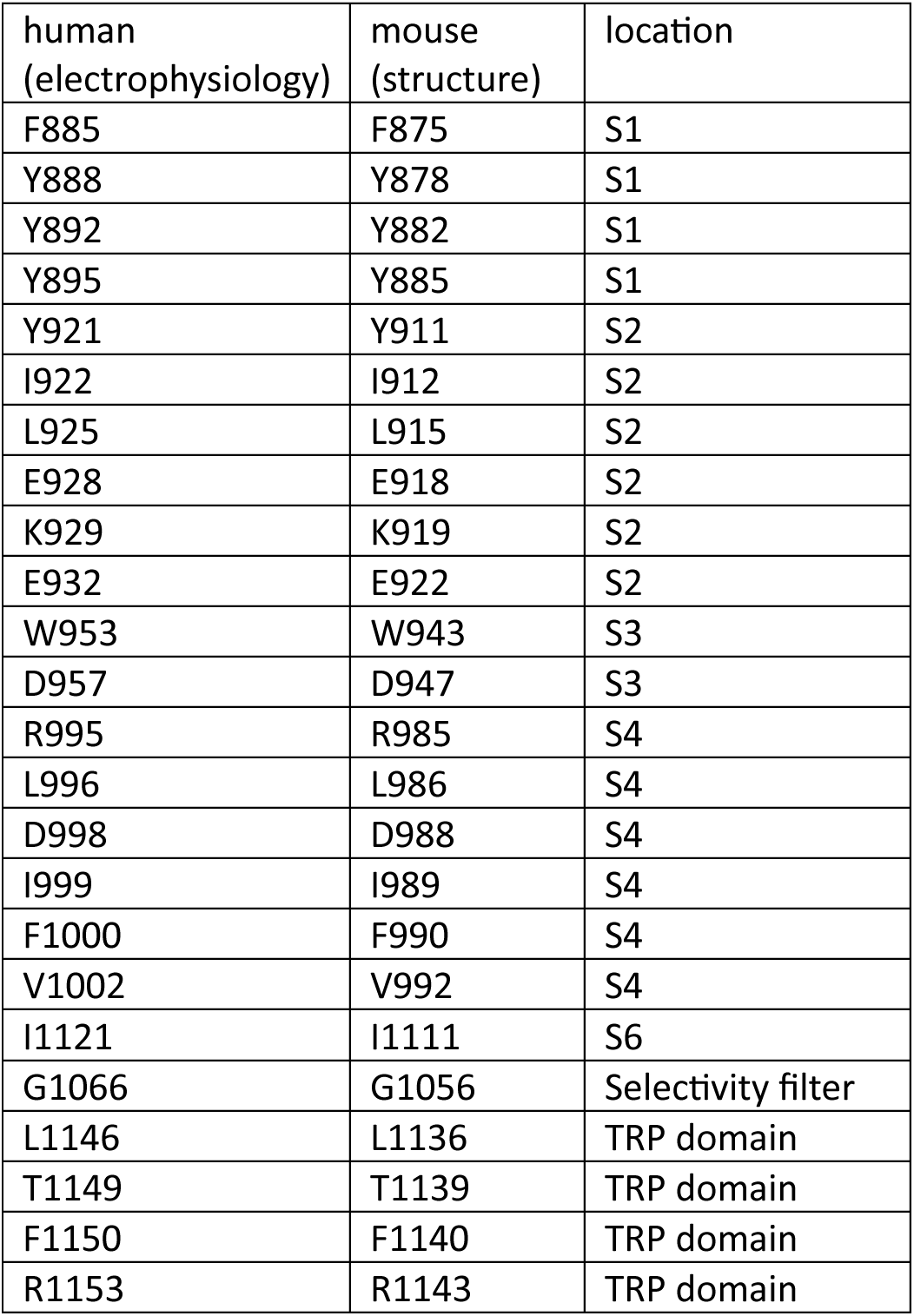
Numbering and location of residues in human and murine TRPM3a2.

| human<br>(electrophysiology) | mouse<br>(structure) | location |
| --- | --- | --- |
| F885 | F875 | S1 |
| Y888 | Y878 | S1 |
| Y892 | Y882 | S1 |
| Y895 | Y885 | S1 |
| Y921 | Y911 | S2 |
| I922 | I912 | S2 |
| L925 | L915 | S2 |
| E928 | E918 | S2 |
| K929 | K919 | S2 |
| E932 | E922 | S2 |
| W953 | W943 | S3 |
| D957 | D947 | S3 |
| R995 | R985 | S4 |
| L996 | L986 | S4 |
| D998 | D988 | S4 |
| I999 | I989 | S4 |
| F1000 | F990 | S4 |
| V1002 | V992 | S4 |
| I1121 | I1111 | S6 |
| G1066 | G1056 | Selectivity filter |
| L1146 | L1136 | TRP domain |
| T1149 | T1139 | TRP domain |
| F1150 | F1140 | TRP domain |
| R1153 | R1143 | TRP domain |

**Supplementary Table 3.**
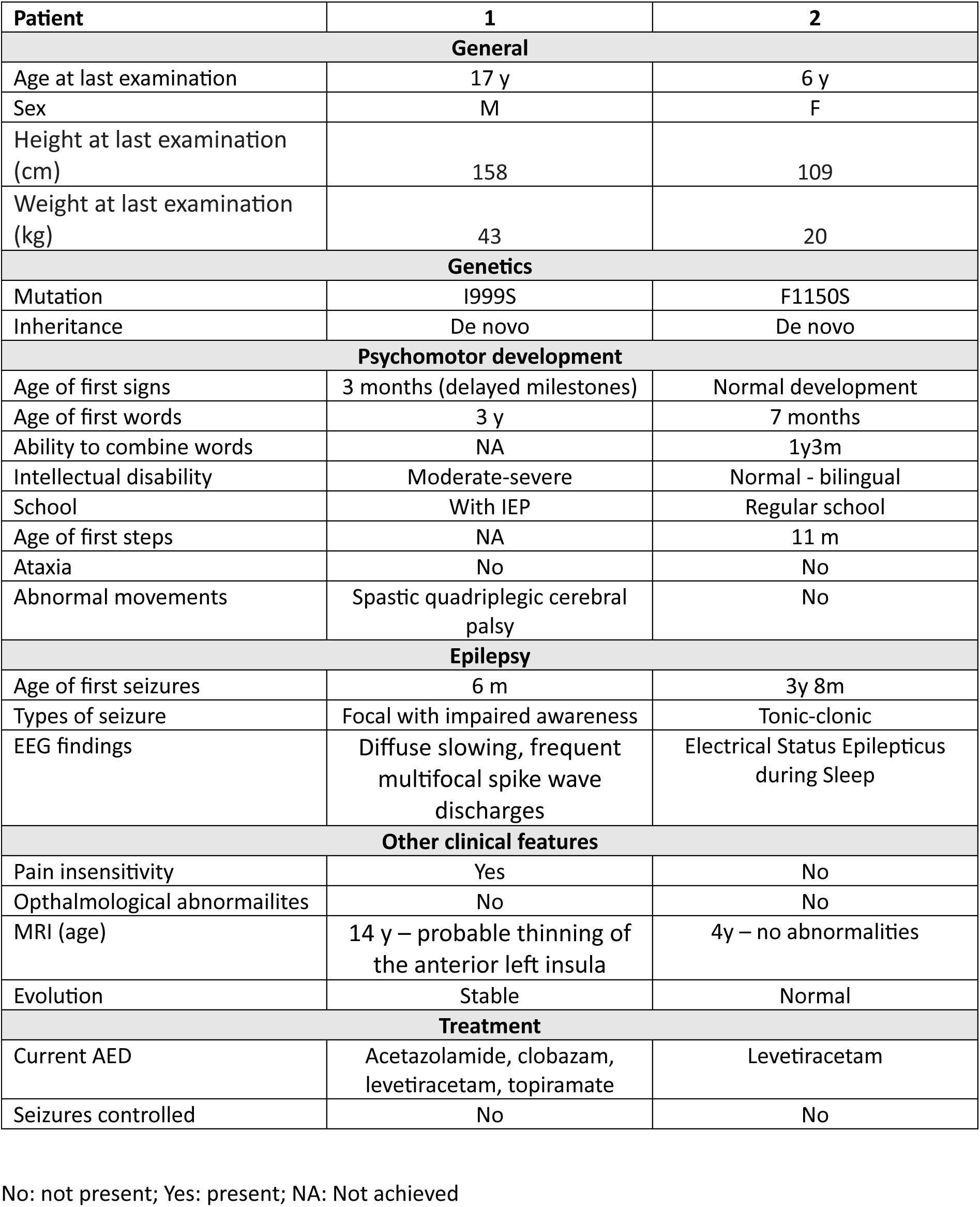
Patient data for I999S and F1150S.

| Patient | 1 | 2 |
| --- | --- | --- |
| **General** |  |  |
| Age at last examination | 17 y | 6 y |
| Sex | M | F |
| Height at last examination (cm) | 158 | 109 |
| Weight at last examination (kg) | 43 | 20 |
| **Genetics** |  |  |
| Mutation | I999S | F1150S |
| Inheritance | De novo | De novo |
| **Psychomotor development** |  |  |
| Age of first signs | 3 months (delayed milestones) | Normal development |
| Age of first words | 3 y | 7 months |
| Ability to combine words | NA | 1y3m |
| Intellectual disability | Moderate-severe | Normal - bilingual |
| School | With IEP | Regular school |
| Age of first steps | NA | 11 m |
| Ataxia | No | No |
| Abnormal movements | Spastic quadriplegic cerebral palsy | No |
| **Epilepsy** |  |  |
| Age of first seizures | 6 m | 3y 8m |
| Types of seizure | Focal with impaired awareness | Tonic-clonic |
| EEG findings | Diffuse slowing, frequent multifocal spike wave discharges | Electrical Status Epilepticus during Sleep |
| **Other clinical features** |  |  |
| Pain insensitivity | Yes | No |
| Ophthalmological abnormalities | No | No |
| MRI (age) | 14 y – probable thinning of the anterior left insula | 4y – no abnormalities |
| Evolution | Stable | Normal |
| **Treatment** |  |  |
| Current AED | Acetazolamide, clobazam, levetiracetam, topiramate | Levetiracetam |
| Seizures controlled | No | No |

No: not present; Yes: present; NA: Not achieved

